# Molecular basis of DNA recognition by the HMG-box-C1 module of Capicua

**DOI:** 10.1101/2022.03.28.485992

**Authors:** Jonathan Webb, Jeremy J.M. Liew, Andrew D. Gnann, MacKenzie Patterson, Sayantanee Paul, Marta Forés, Gerardo Jiménez, Alexey Veraksa, Daniel P. Dowling

**Affiliations:** Chemistry Department, University of Massachusetts Boston, Boston, MA, USA; Instituto de Biología Molecular de Barcelona-Consejo Superior de Investigaciones Científicas (CSIC), Parc Cientific de Barcelona, 08028 Barcelona, Spain; Institució Catalana de Recerca i Estudis Avançats (ICREA), 08028 Barcelona, Spain; Biology Department, University of Massachusetts Boston, Boston, MA, USA

**Keywords:** CIC, helix-turn-helix (HTH), repressor protein, protein-DNA complex, X-ray crystallography

## Abstract

The HMG-box protein Capicua (CIC) is an evolutionarily conserved transcriptional repressor with key functions in development and disease-associated processes. CIC binds DNA using an exclusive mechanism that requires both its HMG-box and a separate domain called C1, but how these domains cooperate to recognize specific DNA sequences is not known. Here we report the crystal structure of the human CIC HMG-box and C1 domains in complex with an 18-base-pair DNA oligomer containing a consensus octameric CIC binding site. We find that both protein domains adopt independent tri-helical structures that pack against opposite sides of the DNA helix. The C1 domain in particular folds into a helix-turn-helix (HTH) structure that resembles the FF phosphoprotein binding domain. It inserts into the major groove of the DNA and plays a direct role in enhancing both the affinity and sequence specificity of CIC DNA binding. Our results reveal a unique bipartite protein module, ensuring highly specific DNA recognition by CIC, and show how this mechanism is disrupted by cancer mutations affecting either the HMG-box or C1 domains.

## Introduction

The HMG-box protein Capicua (CIC) is a tumor suppressor frequently inactivated in oligodendroglioma, gastric adenocarcinoma and other cancers (Bettegowda *et al*, 2011; Cancer Genome Atlas Research Network, 2014; Icgc Tcga Pan-Cancer Analysis of Whole Genomes Consortium, 2020; Kim *et al*, 2021; Okimoto *et al*, 2017; Simon-Carrasco *et al*, 2017; Tan *et al*, 2018). Originally identified in *Drosophila*, CIC is highly conserved in evolution and controls numerous cellular and developmental processes, acting as a sequence-specific transcriptional repressor (Ajuria *et al*, 2011; Dissanayake *et al*, 2011; Jimenez *et al*, 2000; Jimenez *et al*, 2012; Lee *et al*, 2011; Park *et al*, 2017; Simon-Carrasco *et al*., 2017). It often functions as a default repressor downstream of Receptor Tyrosine Kinase (RTK) signaling pathways, which once activated lead to phosphorylation and inactivation of CIC and, consequently, to derepression of its target genes (Astigarraga *et al*, 2007; Futran *et al*, 2015; Liao *et al*, 2017; Okimoto *et al*., 2017; Paul *et al*, 2020; Simon-Carrasco *et al*, 2018; Tseng *et al*, 2007; Wang *et al*, 2017). This connection to RTK signaling means that oncogenic RTK activation can similarly lead to CIC inactivation and derepression of CIC targets such as the ETV1/4/5 family of proto-oncogenes (Bunda *et al*, 2019; Kawamura-Saito *et al*, 2006; Okimoto *et al*., 2017). Aberrant transcriptional activity of CIC is implicated in other clinical disorders, particularly in Spinocerebellar Ataxia Type 1 and other neurobehavioral syndromes (Fryer *et al*, 2011; Lam *et al*, 2006; Lu *et al*, 2017).

CIC contains an HMG-box domain of the Sox type but appears to employ its own mode of DNA binding (Figure 1a) (Forés *et al*, 2017). Unlike Sox proteins, which bind DNA as dimers or together with partner proteins that recognize adjacent DNA sites (Kamachi & Kondoh, 2013), CIC primarily binds to isolated TGAATGAA-like octameric sites independently of other factors. However, *Drosophila* Cic can also cooperatively bind to suboptimal binding sites together with Dorsal, an NF-κB family transcription factor (Papagianni *et al*, 2018). CIC associates with both types of DNA sites by using two separate domains: the N-terminal HMG-box and a C-terminal C1 domain conserved in all CIC proteins (Forés *et al*., 2017; Papagianni *et al*., 2018); how this binding occurs, however, is not known. The importance of the HMG-box and C1 domains for CIC DNA binding is highlighted by the fact that they are both hotspots for inactivating mutations in oligodendroglioma and other tumors (Figure 1b) (Bettegowda *et al*., 2011; Gleize *et al*, 2015). Also, both domains appear to be required for the activity of oncogenic CIC-DUX4 fusions that bind to and activate (rather than repress) CIC targets and cause Ewing-like sarcomas (Forés *et al*., 2017; Graham *et al*, 2012; Kawamura-Saito *et al*., 2006). Recently, rapid release of CIC from DNA has been suggested as a likely first step involved in CIC downregulation by RTK signaling, although the molecular details of this process are lacking (Keenan *et al*, 2020). Therefore, structural studies identifying the mechanism of HMG-box-C1 DNA binding are needed; these studies will advance our understanding of CIC function and regulation and may offer future avenues for blocking CIC-DUX4 activity.

**Figure 1.**
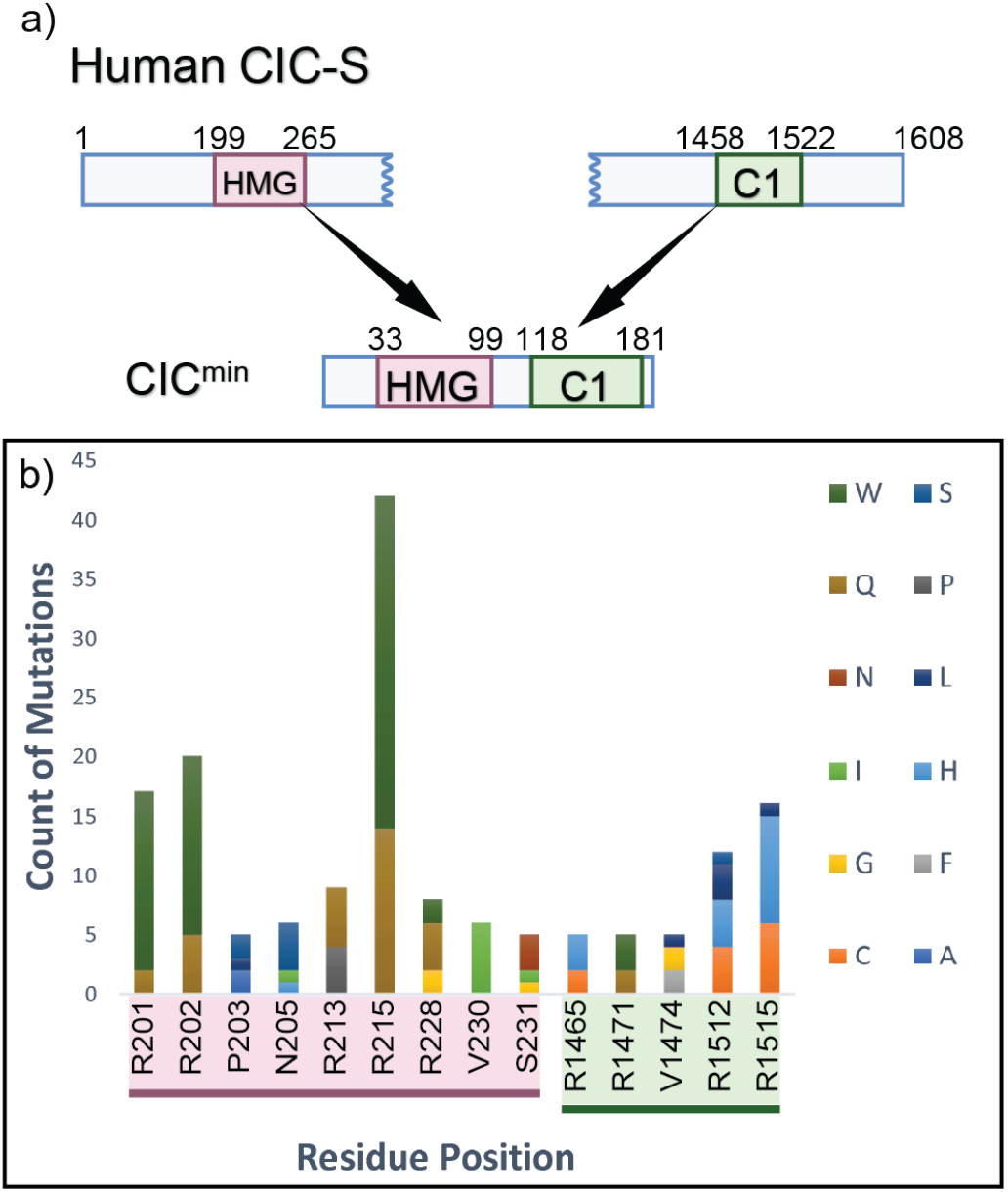
Capicua is a transcription repressor found to contain multiple cancer-associated mutations within its DNA-binding domains. **a)** Shown is a diagram of the CIC-S isoform displaying its two DNA-binding domains, the HMG-box domain (residues 201 – 269) and C1 domain (residues 1459 – 1521). This work employed a minimal construct (CIC^min^) containing the HMG-box and C1 domains joined by an 18-mer linker. **b)** Mutations associated with cancer patients from the COSMIC database were identified within CIC, and those mutations that lie within the CIC DNA-binding domains are displayed. Displayed data have been culled to show only residues that contain at least 5 reported occurrences of a missense mutation.

Here, we explore the structural features of how CIC binds DNA through its HMG-box and C1 domains. We report the crystal structure of the human CIC HMG-box fused to the C1 domain by a short protein linker (an HMG-box-C1 module), in complex with a DNA site from the *ETV5* promoter. We find that the C1 domain adopts a helix-turn-helix (HTH) structure that is necessary for increasing both the affinity and sequence specificity of the HMG-box towards the target site. Additionally, molecular dynamics (MD) simulations were employed to probe fluctuations of the HMG-box and HTH domains when bound to DNA. To our knowledge, this study provides the first molecular depiction of how the HMG-box can be coupled with an HTH domain to bind a highly invariant octameric sequence in DNA.

## Results and Discussion

### The overall structure of the HMG-box and C1 domains of CIC in complex with DNA

To structurally characterize the DNA-binding domains of CIC, an expression construct (CIC^min^) including the HMG-box and C1 domains of the human CIC protein joined by an 18-residue linker was employed (Figure 1a and S1). The decreased size of the expression construct (186 residues) in comparison to full length CIC (1,608 residues in the smaller isoform) was predicted to facilitate crystallization of the protein-DNA complex. We first assessed the ability of CIC^min^ to bind an optimal TGAATGAA sequence by performing electrophoretic mobility shift assay (EMSA) experiments with a labeled DNA probe. As shown in Figure S2, the DNA probe shows the expected mobility shift for a protein-DNA complex when incubated with CIC^min^ at 100:1 ratio of protein:DNA (Forés *et al*., 2017).

The 2.95-Å resolution crystal structure of CIC^min^ in complex with an 18-mer DNA oligonucleotide was solved by molecular replacement using a model of the HMG-box with DNA. The crystal contains one CIC^min^-DNA complex per asymmetric unit in space group *P*2_1_2_1_2_1_ with a Matthew’s coefficient of 2.84 Å^3^/Da. Clear electron density is observed for the HMG-box and C1 domains, as well as the 18-mer DNA (Figure 2a and S3), whereas the N- and C-termini and the interdomain linker are disordered within the crystal structure. The final model includes the HMG-box (His33–Lys106 in PDB 7M5W | His199–Lys272 in Capicua (GenBank ID AAK73515.1)) and C1 (Pro118–Ala180 in PDB 7M5W | Pro1459–Ala1521 in Capicua) domains and the entire 18-mer duplex DNA. Here we will refer to the Capicua numbering (see Figure S4 for sequence numbering). The HMG-box and C1 domain orientations were tethered with a generated linker using BioLuminate (Schrodinger), further validating the complex of CIC^min^ bound to one TGAATGAA sequence.

**Figure 2.**
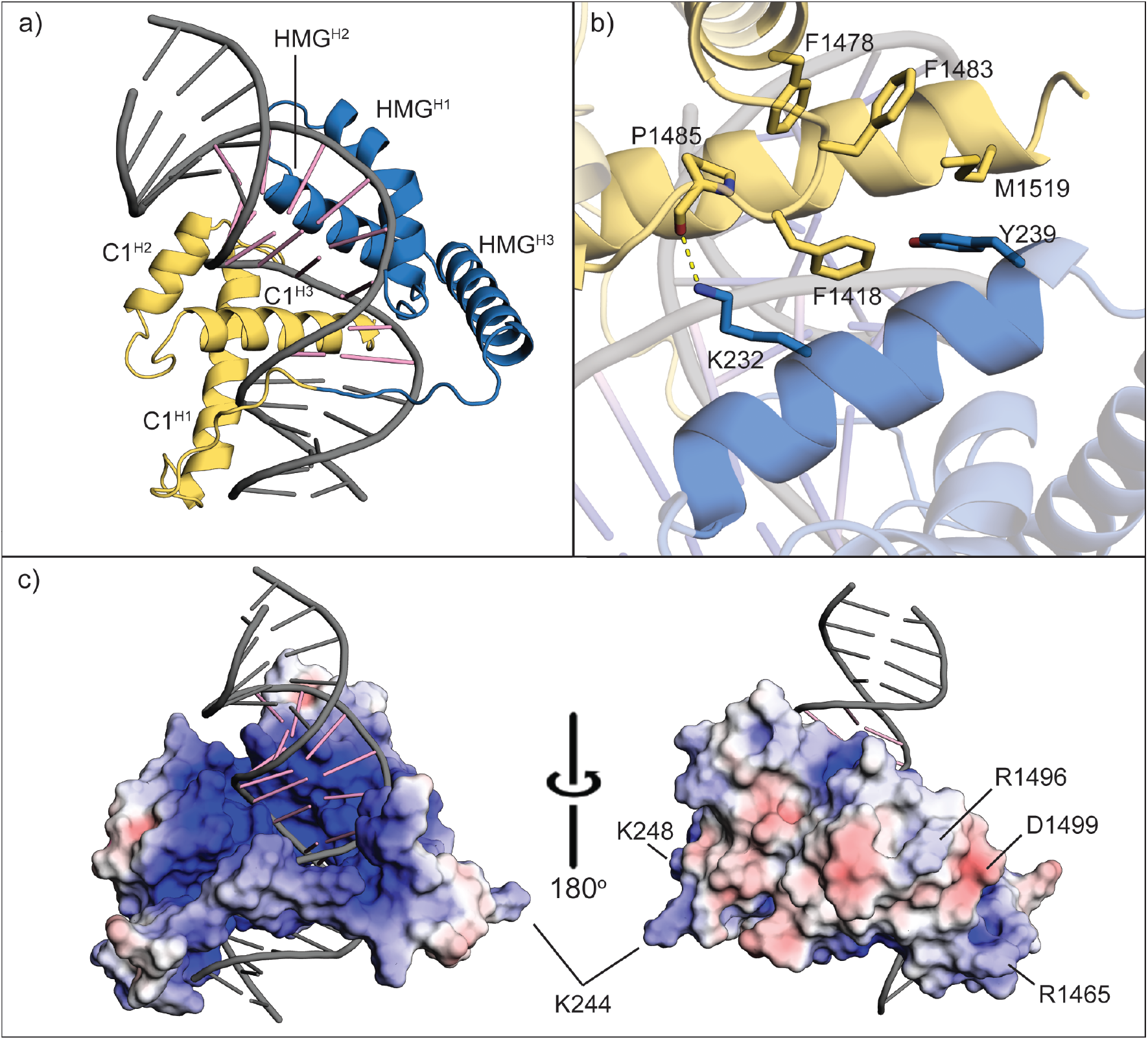
CIC^min^ utilizes coordinated binding of the HMG-box and C1 domains to the core octameric DNA sequence. The C1-domain is colored in yellow, the HMG-box is colored in blue, and the DNA is labeled in pink for nucleotides of the octameric TGAATGAA sequence and grey for bases outside the octameric sequence. **a)** The overall structure of the CIC^min^ construct shows bending of bound DNA. **b)** Shown are the interfacing residues between the HMG-box and C1 domains. **c)** The electrostatic interface of CIC^min^ with DNA reveals an electropositive region of CIC for the binding of DNA. The electrostatic surface is colored as a red to blue heat map for -5 to +5 kBT/ec.

The CIC HMG-box domain adopts a canonical HMG-box fold consisting of three α-helices arranged in an L-shaped configuration, interacting with the minor groove of the DNA along the octameric sequence (Figure 2a). Helix-H1 (HMG^H1^) is inserted within the minor groove of DNA, helix-H2 (HMG^H2^) packs along the DNA phosphodiester backbone, and helix-H3 (HMG^H3^) makes interactions with both the phosphodiester backbone and the minor groove. Binding of the HMG-box induces a ∼66° bent conformation of the DNA calculated using Curves+ (Blanchet *et al*, 2011). The nearest structural homologs are the Sox9 (PDB 4S2Q) (Vivekanandan *et al*, 2015) and Sox18 (PDB 4Y60) (Klaus *et al*, 2016) structures, with root mean square deviations (r.m.s.d.) of ∼1.0 Å for 70 aligned Cα atoms and shared sequence identities of 36% and 39%, respectively (Table S2) (Holm, 2020).

The C1 domain, on the other hand, shares no sequence identity with known protein structures. To our surprise, electron density for the C1 domain revealed a tri-helical HTH fold with helices arranged in a right-handed helical bundle (Figure 2a). In comparison to traditional HTH proteins (Aravind *et al*, 2005), an extended loop is observed between the first and second helices, and a short 3_10_ helix is observed within the loop segment between the second and third helices. At the center of the helical bundle is a hydrophobic core of nonpolar and aromatic residues. The first half of helix-H1 (C1^H1^) is positioned to interact with the DNA phosphodiester backbone. Helix-H2 (C1^H2^) is in an antiparallel orientation to C1^H1^, leading to helix-H3 (C1^H3^), which is inserted into the major groove of the DNA. The major groove where C1 is bound is distorted as a result of HMG-box binding and appears narrower than standard B-form DNA. The classic HTH domain is known to bind to B-form DNA (Aravind *et al*., 2005), and to our knowledge, our structure of the CIC-C1 domain represents the first example of a HTH domain bound to bent DNA.

The orientation of the C1 domain positions the loop preceding C1^H2^ and the C-terminal portion of C1^H3^ closest to HMG^H2^ (Figure 2b). This orientation of domains places HMG-Y239 against C1-M1519 – F1483 – F1484 – F1478 and is stabilized via hydrophobic packing that extends to the hydrophobic core of C1 and additional packing interactions with the DNA. HMG-K232 is positioned within van der Waals distance to the C1 domain as well as hydrogen bonding distance to the backbone C=O group of C1-P1485. The number of interactions between the HMG-box and C1 domains are nevertheless small, which is consistent with the requirement for a covalent linker between the HMG-box and C1 domains for DNA binding (Forés *et al*., 2017). The HMG-box domain aligns with an r.m.s.d. of 0.5 Å for all Cα atoms compared to the recently reported structure of the CIC HMG-box alone with DNA (Lee & Song, 2019), suggesting that despite the observed interactions between the HMG-box and C1 domains, the presence of the C1 domain does not greatly impact the conformation of the HMG-box domain.

The C1 domain is structurally similar to other HTH domains, including the FF domain (a type of HTH domain characterized by two conserved Phe residues located in the first and third helices) of the glucocorticoid receptor DNA-binding factor 1 (PDB 2K85) (Bonet *et al*, 2009), with an r.m.s.d. of 2.3 Å for 60 aligned Cα atoms despite only 12% sequence identity (Figure S5) (Bedford & Leder, 1999). A similar alignment score is observed for other HTH family members including the homeodomain (Table S2). The FF domain, however, contains an inserted 3_10_ helix between the second and third helices, similar to the C1 domain. Interestingly, the FF domain binds phosphopeptides (Aravind *et al*., 2005), and the FF domain from CA150 has been implicated in DNA binding (Lu *et al*, 2009). No structures of the FF domain bound to DNA or to a phosphopeptide are currently available; however, NMR titration analyses of the FF domain from human HYPA/FBP11 support a conserved role of this domain in phosphopeptide binding (Allen *et al*, 2002). Since CIC repressor activity is inhibited through RTK-induced phosphorylation, it is tempting to consider if the structural similarity of the C1 and FF domains might implicate C1 in recognizing a phosphopeptide. Residues within CIC^min^ at the DNA-binding interface create a complementary electrostatic surface for DNA binding (Figure 2c); however, an electrostatic potential mapping of the C1 domain shows the surface away from the DNA is largely nonpolar, except for the presence of R1465 from H1, R1496 of H2, and D1499 within the 3_10_-helix (Lee, 2020). The lack of positive electrostatic features outside of the DNA-binding surface within C1 is inconsistent with C1 solely binding a phosphorylated peptide while bound to DNA, and the surface electrostatic features of the HMG-box domain similarly lack extensive positive electrostatic features outside of a minor patch near R213 of HMG^H1^ and K244 and K248 of HMG^H3^ (Figure 2c).

### The HMG-box and C1 domains of CIC provide a stable DNA-binding interface

The observed binding mode of the HMG-box and C1 domains to bent DNA with a narrower major groove suggests binding of each domain is necessary to increase the binding affinity of the other domain. In fact, neither domain can bind without the other under physiological conditions (Forés *et al*., 2017), which suggests that the intact CIC protein must maintain functional orientations of both the HMG-box and C1 domains to achieve robust DNA binding. To explore the stability of the CIC^min^-DNA complex, motions of the complex on the ns time scale were calculated from three separate 250 ns runs of MD simulation within Schrodinger’s BioLuminate. As expected, the root mean square fluctuation (r.m.s.f.) values of loops in both domains are greater than the ordered α-helices, and the protein termini and interdomain linker have the highest r.m.s.f. values (Figures 3a and S6). Within the C1 domain, helix H3 demonstrates the lowest r.m.s.f. values, consistent with its placement towards the narrow major groove, at the interface with the phosphodiester backbone of the lower strand. A diagram showing CIC^min^-DNA interactions that are stable over the course of the 250 ns trajectories is displayed in Figure 3b.

**Figure 3.**
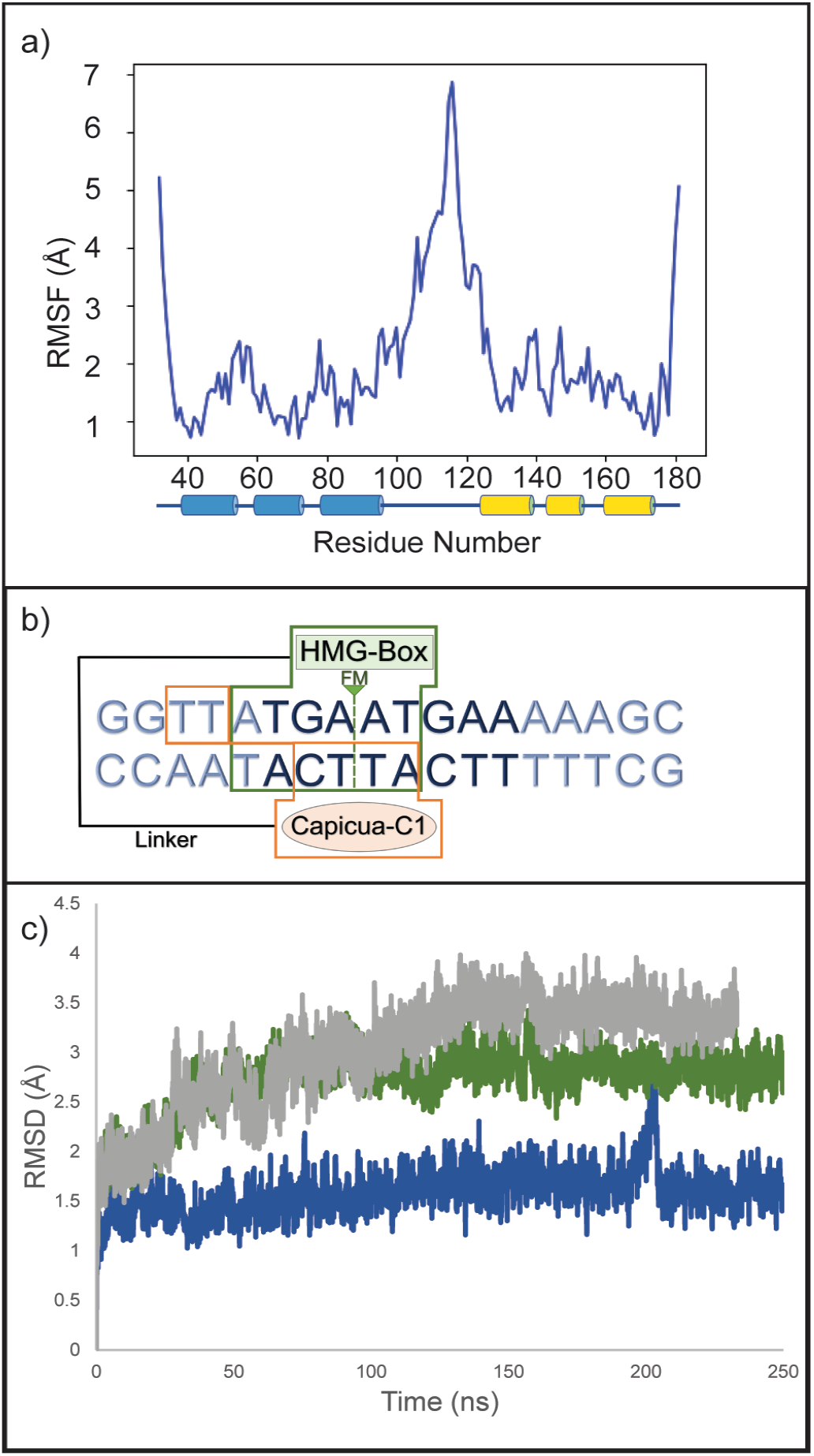
Molecular dynamics simulations of the CIC^min^ structure show stable binding of the DNA oligomer. **a)** The r.m.s.f. values calculated over a representative 250 ns trajectory shows the linker region as being highly flexible relative to the HMG-box and C1 domains (represented by blue and green cylinders for helices, respectively). See Figure S7 for information for all MD simulations. **b)** Contacts between CIC^min^ and the DNA oligomer over the trajectory reveal interactions between the C1-domain and DNA (orange boxing) and the HMG-box with DNA (green boxing) that are present in over 50% of the MD simulation. **c)** The r.m.s.d. values, calculated from the alignment Cα of protein helices to the initial frame for the three trajectories, indicates equilibration of the CIC^min^-DNA complex with DNA binding preserved as indicated in a latter trajectory timepoint (see Fig. S7).

We further used MD simulations to examine the influence of the interdomain linker on the CIC^min^-DNA structure. The r.m.s.d. values of helices of the HMG-box and C1-domain over the 250 ns trajectories are shown in Figure 3c. The structures appear to equilibrate within the first half of the trajectories, and an alignment of initial and final states shows similar positioning of the HMG-box and C1-domains (Figure S7 and S8a). Interestingly, the linker behaves differently in the three performed simulations, suggesting it does not pose large restraints on DNA binding (Figure S8b). Further inspection confirms that the linker is long enough to accommodate movement of the C1 domain away from the HMG-box along the DNA (Figure S9). The helices surrounding the linker, HMG^H3^ and C1^H1^, are more mobile over the course of the trajectories and have higher r.m.s.f. values compared to helices that have greater contacts with the DNA (Figures S6 and S7). Therefore, the CIC^min^-complex structure is likely unaffected by the linker and faithfully represents interactions between the C1 domain, the HMG-box domain, and DNA. This suggests that binding of CIC to DNA is similarly independent of sequences located outside the HMG-box and C1 domains, with both domains freely moving and aligning during binding.

### Molecular determinants of CIC^min^ binding to DNA

The first and second helices of the HMG-box and the N-terminal residues before the first helix make the majority of interactions between the HMG-box and the DNA. HMG^H1^ bends the oligomer DNA by packing F207 and M208 as a wedge between subsequent DNA bases in the minor groove of the octameric sequence (Figure 4a). Two residues N-terminal to the HMG^H1^ helix, R201 and R202, wrap around the DNA to enable possible hydrogen bond and electrostatic contacts (Figure 4b). R201 is directed towards phosphates of the backbone while R202 is positioned more closely to the first dT and dG of the octameric sequence. Within HMG^H1^, K212 interacts with the phosphodiester backbone whereas R215 is positioned to interact with dT10 of the DNA as well as N227 and S231 of HMG^H2^ (Figure 4c). Within the turn between the HMG^H1^ and HMG^H2^ helices, D226 can form a water-mediated interaction with dA12 in the octameric sequence and N227 can form hydrogen bond contacts with dT10 and dG11 of the octameric sequence. S231 forms a hydrogen bonding interaction with the N3 nitrogen of the second adenine in the lower strand of the CIC site. By contrast, HMG^H3^ seems to play a minor role in binding, with only a few contacts to the DNA backbone. H250 contributes to the interface facing the oligomer along with K257 and N205 (from the N-terminus), which form two hydrogen bonding contacts with the phosphodiester backbone of the lower strand (Figure 4d).

**Figure 4.**
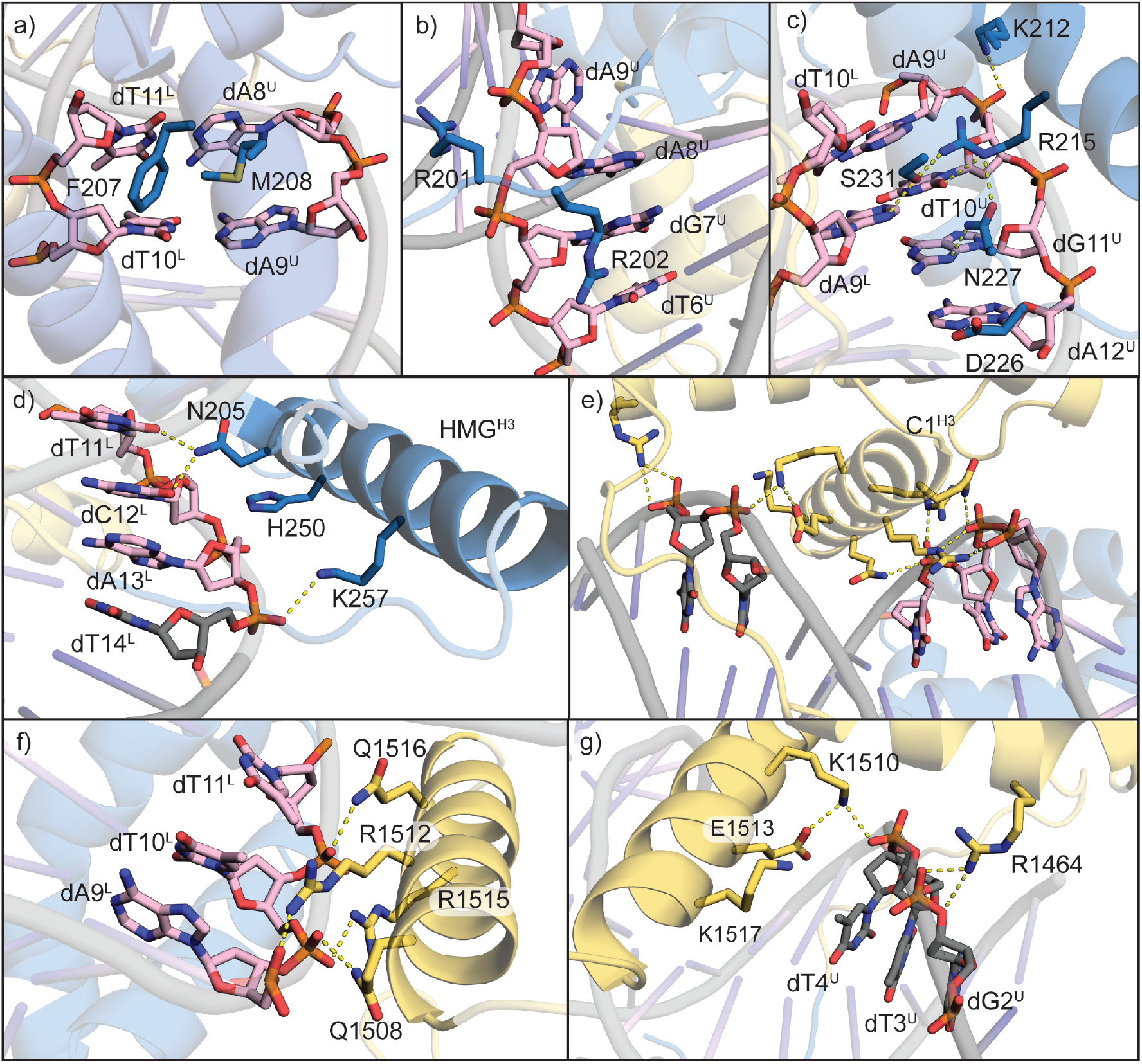
The crystal structure of CIC^min^ details specific interactions by the HMG-box and C1 domains within and around the octameric sequence of the DNA. The first four panels highlight interactions of the HMG-box (blue) with DNA and the final three panels show the interactions between residues of helix C1^H3^, of the C1 domain (yellow), with DNA colored pink for nucleotides of the octameric sequence and grey for all other nucleotides. Superscripts indicate upper (U) or lower (L) DNA strand. Interactions within 3.3 Å are shown as yellow dashed lines. **a)** Residues F207 and M208 intercalate between two AT pairs of DNA in the octameric sequence. **b)** Residues R35 and R36 wrap around the phosphodiester backbone of the first TGAA DNA sequence. **c)** The first two helices of the HMG-box domain donate several polar interactions with the upper strand just after the intercalation by F207 and M208. **d)** Helix HMG^H3^ forms several interactions with the backbone of the DNA. **e)** The C1 domain aligns the third helix within the major groove of the DNA, where it forms interactions with the DNA backbone **f)** within and **g)** outside of the octameric recognition sequence. Panels f and g are rotated approximately 180° about a plane vertical to panel e.

Our structure suggests why the HMG-box alone does not allow for binding of DNA oligomer by CIC under physiological conditions. In the CIC^min^ structure, the C1 domain makes several non-sequence specific contacts that may increase binding affinity of CIC to DNA. The majority of these contacts originate from the C1^H3^ helix and are exclusively with the phosphodiester backbone within the major groove, opposite of the HMG-box (Figure 4e). These interactions are within the complement octameric sequence as well as outside of the recognized octameric sequence. Q1516, R1512, R1515, and Q1508 are within hydrogen-bonding distance of 5’-A^9^-T^11^-3’ of the complement octameric sequence (Figure 4f). The C1^H3^ helix also forms contacts with bases upstream of the consensus octameric sequence. K1510 and R1461, from the C1^H1^ helix, can form hydrogen bonds with the phosphates of these nucleotides (Figure 4g). K1510 is also within distance to form a hydrogen-bonding network with E1513. These interactions likely support an important role of C1^H3^ in contacting the lower strand of the CIC site via the major groove.

### DNA specificity of CIC is driven by an unpredicted DNA binding mode for a HTH domain

Clear electron density enabled unambiguously building the entire 18-mer duplex DNA. Supporting the DNA directionality in our model, refinement of the DNA in the flipped orientation led to an increase in R-free of ∼2.6% and difference electron density peaks (Figure S10a and S10b). The orientation of CIC in our structure positions the intercalating residues of the HMG-box domain, M208 and F207, between the 5’-A^8^A^9^-3’ of the first half of the CIC octameric sequence and 3’-T^10^T^11^-5’ of the complement, respectively. Surprisingly, in the previously reported structure of the CIC HMG-box alone with DNA (PDB ID 6JRP) (Lee & Song, 2019) the octameric sequence is flipped to interact with the opposite strand, where the HMG-box disrupts base stacking with F207 between the second 5’-GA-3’ of the octameric sequence and M208 between the complement 3’-CT-5’ (Figure S11). Since our previous studies revealed that the HMG-box alone does not effectively interact with DNA (Forés *et al*., 2017), the above differences in binding led us to wonder if the C1 domain plays a role in conferring DNA binding specificity.

To address C1 contributions to DNA specificity, we analyzed movement of the C1 domain over the 250 ns simulations. From the solved structure, the C1 domain is positioned with C1^H3^ along the major groove making most of the contacts with the DNA, and this binding mode is maintained during the trajectory (Figures 3 and 4e and Figure S7). Contacts between the C1 domain and the DNA are strictly with the DNA phosphodiester backbone throughout MD simulations, either by hydrogen bonding with polar residues such as R1464, K1510, R1512, and R1515, or by water-mediated contacts with Q1508, E1513, and Q1516 (Figure 4e - g). E1513 makes a water-mediated contact between its carboxylate and the phosphate group of the second of the thymines. R1512, R1515, Q1508, and Q1516 make contacts with the phosphate groups of the 5’-A^4^T^5^T^6^C^7^-3’ of the complement strand. These C1 interactions lack hydrogen-bonding specificity with the octameric sequence; however, Q1516 is positioned at the second nucleoside of the octamer, and its location near dC7 of the lower strand would clash with a C5 methyl group of a thymidine in this position, as well as with the observed dA in the 6JRP structure through positioning the Q1516 C=O group near the lone pair of N7 of adenine. Therefore, the C1 domain does not seem to provide specificity through specific interactions with the DNA, but through exclusion of alternative interactions. The C1 domain likely provides stability in the bound structure through increased interactions with the DNA and possible interactions with HMG^H2^ along the DNA backbone.

### Cancer mutations within the CIC DNA-binding domains

To better understand the impact of cancer-associated CIC mutations, we used mutational data from the COSMIC database (Tate *et al*, 2019) to evaluate the effects of tumor-validated substitutions on the CIC^min^:DNA structure (Figure 5). The most frequent mutations affecting the HMG-box correspond to arginine residues (R201, R202, and R215) changing to either a bulkier tryptophan residue or to a shorter glutamine residue (Figures 1b, and S12). R201 and R202 are both located at the N-terminal region of the HMG-box, prior to HMG^H1^, where R202 is positioned to hydrogen bond with the C2 position carbonyl of dT6 and the N3 nitrogen of dG7, and R201 is positioned to hydrogen bond with the phosphodiester backbone of dA8 (Figure 4b – 4d). The R215 residue is the most frequently associated somatic mutation within the COSMIC database and is most frequently mutated to either glutamine or tryptophan. R215 makes a number of water-mediated contacts, including to the C2 position carbonyl of dA9 of the complement strand and the oxygen of the ribose dT10 within the upper strand, over the course of the 250 ns trajectory. The mutations of these residues to either glutamine or tryptophan would disrupt those interactions and potentially sterically hinder CIC from binding to its target octameric sequence. Indeed, loss of DNA binding may represent the primary molecular defect underlying CIC-associated cancers (Forés *et al*., 2017).

**Figure 5.**
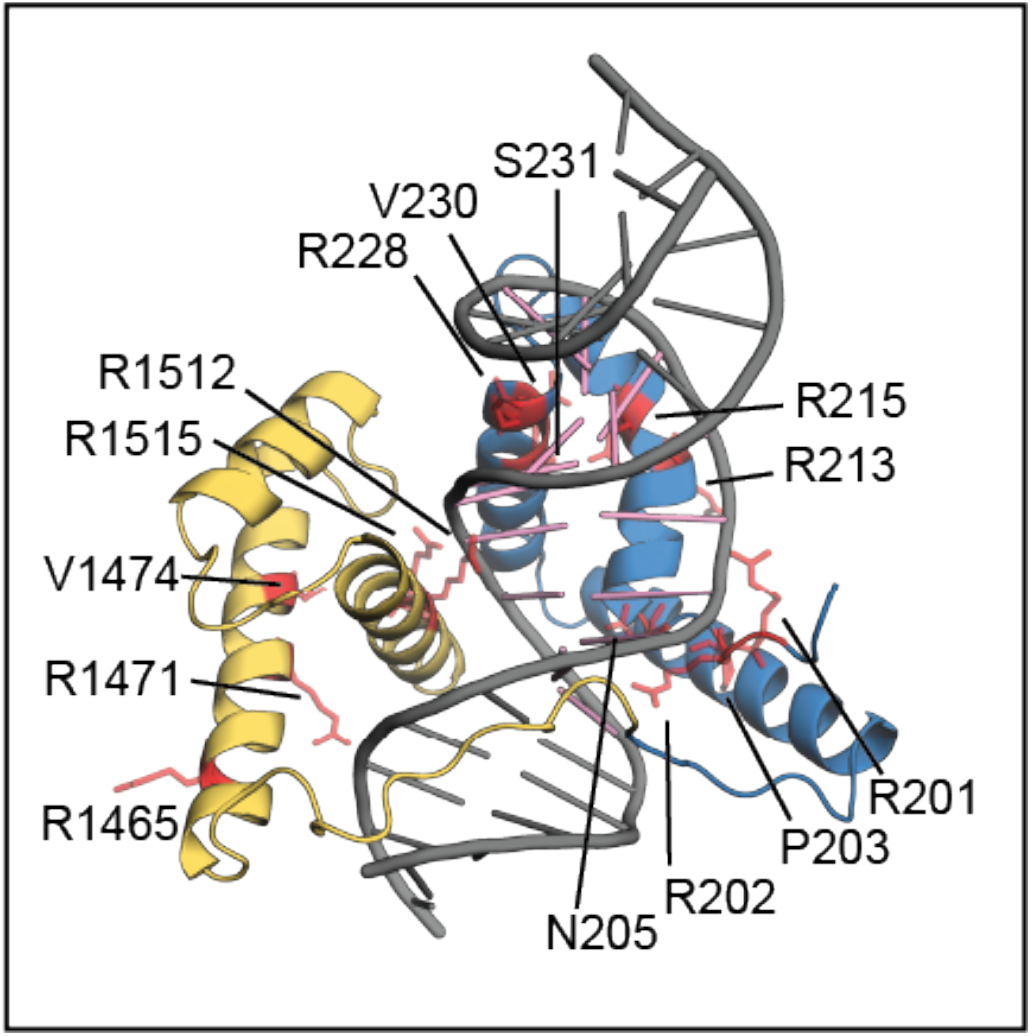
COSMIC mutations within CIC^min^ mapped onto the structure demonstrate residues predicted to be linked to DNA binding and/or protein stability. Domains are colored as in Fig. 2, and COSMIC mutations are colored red and displayed as sticks.

There are also several mutations found in the COSMIC database within the C1 domain. Missense mutations primarily affect R1465, R1471, V1474, R1512, and R1515 (Figure S13). Unlike in the case of the HMG-box, the most frequent mutations are arginine to methionine, cysteine or leucine. V1474 is located on C1^H3^ and is wedged between the first and third helices of the C1 domain; therefore, mutations of V1474 to similarly non-polar residues leucine, glycine, and phenylalanine may alter hydrophobic packing between the helices of the C1 domain and disrupt DNA binding through overall decreased stability. R1512 and R1515 are positioned along the backbone of the DNA, making hydrogen bonding interactions with the phosphodiester backbone of 5’-A^9^T^10^T^11^-3’ of the complement strand. These residues are the most frequently mutated in CIC-associated cancers, and the structure of CIC^min^ with DNA provides a clear rationale; loss of the positive electrostatic interactions and hydrogen bonding capability of either R1512 or R1515 would severely disrupt the protein-DNA interface, thus decreasing CIC DNA binding and regulatory activity.

In summary, we have presented the structure of the CIC HMG-box and C1 bi-partite module bound to DNA, revealing how proper CIC repressor activity and inactivation by cancer mutations are linked to both domains. The C1 domain adopts an HTH fold that resembles the overall structure of the FF domain, uncovering a previously unknown mechanism of DNA binding in which an HMG-box and an HTH domain act together to recognize a specific octameric target site. Interestingly, while the HMG-box is an ancient domain present in a diverse set of proteins, the C1 domain is restricted to CIC family proteins, suggesting that the latter originated through mutation and selection within a pre-existing HMG-box protein. This would have allowed a highly selective mechanism of recognition that is not found among other HMG-box factors, but which is nevertheless susceptible to inactivation by mutations affecting either of those two domains, as seen in CIC-related cancers. Finally, since oncogenic CIC-DUX chimeras rely on the same DNA binding mechanism for activating instead of repressing CIC target genes, the structure we report here may facilitate the development of strategies to interfere with CIC-DUX activity in CIC-rearranged sarcomas.

## Materials and Methods

### Materials

Cell lines and molecular biology materials were purchased from New England Biolabs (NEB) unless otherwise indicated. Crystallization reagents were purchased from Hampton Research. Other chemicals were purchased from commercial suppliers and used without further purification.

### Protein overexpression and purification

A minimal DNA-binding construct (HMG-box-C1, termed CIC^min^) containing the HMG-box (D190 to N273) and C1 (K1459 to Q1529) domains of human Capicua (GenBank ID AAK73515.1) was generated with an 18-amino acid linker containing 15 residues flanking helix H3 of the HMG-box, a phenylalanine insertion for increased UV absorption to aid protein purification, and two residues that precede H1 of the C1 domain. CIC^min^ was generated by GenScript as a codon-optimized gene sequence for recombinant protein expression in *E. coli* (Figure S1). The commercially synthesized DNA product was digested with NdeI and BamHI in CutSmart Buffer and purified using a Monarch PCR and DNA Cleanup Kit. The pET28a vector was similarly digested, treated with alkaline phosphatase, and purified on a 1.0% (w/v) agarose gel. The digested vector was extracted using the Monarch DNA Gel Extraction Kit, and the digested CIC^min^ DNA and pET28a were ligated together in a 5:1 ratio using T4 DNA ligase, yielding pET28^cicmin^. Ligated product was used to transform NEB5α cells. A single colony was grown overnight in 5 mL Miller lysogeny (LB-Miller) medium supplemented with 50 µg/mL kanamycin sulfate. DNA was isolated using a Monarch Plasmid Miniprep Kit, and Sanger DNA sequencing was performed by Genewiz.

Chemically competent T7 Express cells were transformed with pET28^cicmin^ and plated on LB-Miller agar supplemented with 50 µg/mL kanamycin. A starter culture originating from a single colony was used to inoculate 8 × 1 L of autoclaved LB-medium with 50 µg/ml kanamycin and grown at 37 °C while shaking at 250 rpm until an optical density (OD_600_) of 0.50 was reached. The culture was induced by adding IPTG to a concentration of 0.1 mM and the cells were grown for 4 h at 37 °C shaking at 250 rpm. Cells were harvested via centrifugation at 4,816 × g for 30 min at 4 °C, and the resulting cell pellet was resuspended in Buffer A (50 mM Tris (pH 8.0), 500 mM sodium chloride, and 10% (v/v) glycerol) and supplemented with an EDTA-free Pierce Protease Inhibitor tablet. The resuspended cells were kept on ice and lysed by sonication using a Branson Digital 250 Sonifier with five cycles of 3 min active sonics (40% duty, output level 7) and 3 min rest. Cell debris was removed by centrifugation at 20,000 × g and 4 °C for 1 h. The supernatant was consecutively filtered through 0.45 µm and 0.22 µm PES syringe filters prior to loading onto a 5 ml HisTrap nickel affinity column (GE Healthcare) preequilibrated with Buffer A. Unbound protein was removed using an isocratic step with 8% Buffer B (50 mM Tris (pH 8.0), 500 mM sodium chloride, 10% (v/v) glycerol, and 1 M imidazole), and bound CIC^min^ was then eluted with a linear gradient from 8 to 50% Buffer B. Fractions containing CIC^min^ were collected and dialyzed using a slide-a-lyzer mini dialysis cassette (3500 MWCO) (Pierce) into Buffer A supplemented with 1 mM DTT. The nickel affinity chromatography step was repeated once more with dialyzed CIC^min^ prior to purification on a HiLoad Superdex 75 16/600 column (GE Healthcare) equilibrated with 50 mM Tris (pH 8.0), 300 mM sodium chloride, 10% (v/v) glycerol, and 1 mM DTT. The size and purity of the eluted protein was determined by SDS–PAGE, and protein was concentrated using a 3 kDa MWCO Vivaspin centrifugal concentrator (MilliporeSigma). Final protein concentration was assessed using 5,5-dithio-bis-(2-nitrobenzoic acid) (ThermoFisher) and a standard curve of glutathione (Aitken & Learmonth, 2003; Winther & Thorpe, 2014). Protein was aliquoted, flash frozen in liquid nitrogen, and stored at –80 °C.

### Electrophoretic mobility shift assay (EMSA)

Fluorescent electrophoretic mobility shift assay was performed using the CIC^min^ protein, expressed and purified as described above. Oligonucleotides were labeled with IRDye 700 (LI-COR). The sequence with a CIC binding site was derived from the *Drosophila huckebein* (*hkb*) promoter (5’-GTCCCAGTTTATGAATGAATTTACTAAATC-3’). Oligonucleotides were diluted in 1× TE to the final concentration of 20 pmol/μL. For annealing, 10 μL of forward and 10 μL of reverse oligonucleotide were mixed in a microcentrifuge tube, placed into a beaker with boiling water, and allowed to cool to room temperature. Annealed oligonucleotides were further diluted to a working concentration of 0.1 pmol/μL. Binding reactions were carried out in a total volume of 20 μL containing 10 mM Tris-HCl (pH 7.5), 50 mM KCl, 3.5 mM DTT, 0.25% Tween 20, 1 μg poly(dI·dC), 0.1 pmol of annealed oligo, and different molar concentrations of the protein. Reactions were incubated for 30 min at room temperature in the dark. Samples were mixed with 1 μL of 10× Orange loading dye (LI-COR) and loaded on a 4% 1× TBE native polyacrylamide gel. The gel was run in the dark for 30 min and scanned with the LI-COR Odyssey imaging system.

### Crystallization and data collection

To obtain a protein-DNA complex, an 18-mer DNA oligonucleotide containing the ETV5 promoter (GGTTA<u>TGAATGAA</u>AAACC) and its reverse complement were purchased from ThermoFisher with HPLC purification. DNA was resuspended in ultrapure water and quantified by absorbance at 260 nm using a NanoDrop 2000c spectrophotometer. CIC^min^ in complex with the 18-mer DNA oligonucleotide was screened for crystallization conditions using a Crystal Phoenix robot (Art Robbins Instruments) within the MIT crystallization facility. A 3:1 ratio of protein:DNA using a final concentration of 7.1 mg/mL CIC^min^ was used in sparse matrix screening, and crystallization conditions were identified using the sitting drop vapor diffusion method with the Kerafast Protein-Nucleic Acid Complex Crystal Screen. Crystals appeared under multiple conditions, and diffraction quality crystals leading to structure determination were obtained in 16% (w/v) PEG 8000, 0.1 M 2-morpholinoethanesulofnic acid (MES) (pH 6.0), 0.1 M CaCl_2_, and 0.1 M NaCl. Crystals were cryoprotected with the reservoir solution supplemented with final concentrations of 20% (v/v) glycerol and 20% (w/v) PEG 8000 and cryocooled in liquid nitrogen.

### Structure determination and refinement

X-ray diffraction data were collected at a wavelength of 1.54178 Å at the MIT crystallization facility on a rotating copper anode X-ray generator (Micromax 007-HF) equipped with a Saturn 944+ detector and cryostream 800 (Oxford Cryosystems). Data were collected with 0.5° oscillations at 100 K. Diffraction intensities were indexed to space group *P*2_1_2_1_2_1_, integrated, and scaled with the XDS program suite (Kabsch, 2010). Data processing details are presented in Table S2.

The structure of the CIC^min^–DNA complex was solved by molecular replacement (MR) using chain A of PDB ID 6JRP as a search probe (Lee & Song, 2019). MR was performed in PHASER (McCoy *et al*, 2007) within Phenix (Liebschner *et al*, 2019), yielding a solution with overall LLG and TFZ scores of 445 and 20.8, respectively. Clear electron density for the full HMG-box domain was observed with remaining difference electron density for the 18-mer DNA and the C1 domain. Model building was performed in COOT (Casañal *et al*, 2020), and refinement was carried out in Phenix (Liebschner *et al*., 2019). Initial refinement steps included simulated annealing with minimization and individual ADP refinement. Subsequent refinement steps included minimization, individual ADP refinement, and translation/libration/screw (TLS) motion refinement with separate TLS groups defining the HMG-box, the C1-domain, and each DNA strand. Waters and a calcium ion were added towards the end of refinement. The model was validated using composite omit maps, and Ramachandran angles were assessed using MolProbity (Williams *et al*, 2018). Final model refinement statistics are presented in Table S2. The final model includes the HMG-box (His33–Lys106) and C1 (Pro118– Ala180) domains of the protein and the entire 18-mer duplex DNA.

### Molecular dynamics

MD simulations were performed through the Desmond suite of BioLuminate 2021 (Beard *et al*, 2013; Bowers *et al*, 2006; Salam *et al*, 2014; Zhu *et al*, 2014). Three MD simulations of CIC^min^ complexed with DNA (PDB ID 7M5W) were performed. The structure was preprocessed using the Protein Preparation Wizard tool available in the Biologics suite of BioLuminate 2021 (Sastry *et al*, 2013) to add hydrogens and optimize hydrogen-bonding. The resulting model was refined to decrease potential energy until the heavy-atom r.m.s.d. reached 0.3 Å, at which point the refinement was stopped. The minimized structure was solvated in SPC water and neutralized by the addition of 19 Na^+^ ions. The structure was heated for 2 ps at 300 K and 1.01325 bar using the Nose-Hoover chain thermostat method, Martyna-Tobias-Klein barostat method, and isotropic coupling. After heating, the simulations were conducted for 250 ns using the NPT ensemble. The OPLS4 force field was used for all simulations (Lu *et al*, 2021). The first MD simulation ran for a length of 250 ns with 100 ps frames and the second and third for a length of 250 ns and 233 ns with 25 ps frames.

Analysis of trajectories was done using the Simulation Event Analysis (SEA) module within BioLuminate 2021. For r.m.s.d. calculations, protein selections were limited to helices of the HMG-box and C1-domains. The trajectories were fit to the initial frame of the respective trajectory after the initial equilibration of the MD simulation. SEA was used to generate r.m.s.f. and r.m.s.d. data for the three trajectories.

## Supporting information

Electronic Supporting Information Figures and Tables

## Figure generation

Protein figures were generate using PyMOL (Schrodinger, 2015) and electrostatic surface calculations were performed with the APBS plugin (Dolinsky *et al*, 2004). MD graphs were plotted using BioLuminate (Schrodinger), and multiple sequence alignment figures were generated using Geneious Prime 2022.0.1 (https://www.geneious.com).

## Coordinate deposition

The coordinates and structure factors have been deposited to the protein data bank (www.pdb.org) as PDB ID 7M5W.

## Acknowledgements

The authors kindly acknowledge the MIT Structural Biology Core facility and Staff Scientist Robert Grant. A.V. is funded by the National Institutes of Health (NIH) grant GM141843. G.J. is funded by the Spanish Government (grants BFU2017-87244-P and PID2020-119248GB-I00). This work was funded in part by the University of Massachusetts Boston Office of Global Programs (to D.P.D.).

## Author contributions

J.W., J.J.M.L., A.D.G., M.P., S.P., and M.F. conducted experiments and curated data; J.W., J.J.M.L., A.D.G. and D.P.D. formal analysis; J.W., J.J.M.L., A.D.G., S.P., M.F., G.J., A.V. and D.P.D. validation; G.J. and A.V. funding acquisition; J.W., G.J., A.V., and D.P.D. wrote the manuscript; G.J., A.V., and D.P.D. resources.

## References

Aitken A, Learmonth M (2003) Quantitation and location of disulfide bonds in proteins. Methods Mol Biol 211: 399–410

Ajuria L, Nieva C, Winkler C, Kuo D, Samper N, José Andreu M, Helman A, González-Crespo S, Paroush Z, Courey AJ et al (2011) Capicua DNA binding sites are general response elements for RTK signaling in Drosophila. Development 138: 915–924

Allen M, Friedler A, Schon O, Bycroft M (2002) The structure of an FF domain from human HYPA/FBP11. J Mol Biol 323: 411–416

Aravind L, Anantharaman V, Balaji S, Babu MM, Iyer LM (2005) The many faces of the helix-turn-helix domain: transcription regulation and beyond. FEMS Microbiol Rev 29: 231–262

Astigarraga S, Grossman R, Diaz-Delfin J, Caelles C, Paroush Z, Jimenez G (2007) A MAPK docking site is critical for downregulation of Capicua by Torso and EGFR RTK signaling. Embo Journal 26: 668–677

Beard H, Cholleti A, Pearlman D, Sherman W, Loving KA (2013) Applying physics-based scoring to calculate free energies of binding for single amino acid mutations in protein-protein complexes. PLoS One 8: e82849

Bedford MT, Leder P (1999) The FF domain: a novel motif that often accompanies WW domains. Trends Biochem Sci 24: 264–265

Bettegowda C, Agrawal N, Jiao Y, Sausen M, Wood LD, Hruban RH, Rodriguez FJ, Cahill DP, McLendon R, Riggins G et al (2011) Mutations in CIC and FUBP1 contribute to human oligodendroglioma. Science 333: 1453–1455

Blanchet C, Pasi M, Zakrzewska K, Lavery R (2011) CURVES+ web server for analyzing and visualizing the helical, backbone and groove parameters of nucleic acid structures. Nucleic Acids Research 39: W68–W73

Bonet R, Ruiz L, Aragón E, Martín-Malpartida P, Macias MJ (2009) NMR structural studies on human p190-A RhoGAPFF1 revealed that domain phosphorylation by the PDGF-receptor ? requires its previous unfolding. J Mol Biol 389: 230–237

Bowers KJ, Chow DE, Xu H, Dror RO, Eastwood MP, Gregersen BA, Klepeis JL, Kolossvary I, Moraes MA, Sacerdoti FD et al (2006) Scalable algorithms for molecular dynamics simulations on commodity clusters. Proceedings of the ACM/IEEE Conference on Supercomputing (SC06), Tampa, Florida: 43–43

Bunda S, Heir P, Metcalf J, Li ASC, Agnihotri S, Pusch S, Yasin M, Li M, Burrell K, Mansouri S et al (2019) CIC protein instability contributes to tumorigenesis in glioblastoma. Nat Commun 10: 661

Cancer Genome Atlas Research Network (2014) Comprehensive molecular characterization of gastric adenocarcinoma. Nature 513: 202–209

Casañal A, Lohkamp B, Emsley P (2020) Current developments in Coot for macromolecular model building of electron cryo-microscopy and crystallographic data. Protein Science 29: 1055–1064

Dissanayake K, Toth R, Blakey J, Olsson O, Campbell DG, Prescott A, Mackintosh C (2011) Erk/p90RSK/14-3-3 signalling impacts on expression of PEA3 Ets transcription factors via the transcriptional repressor capicúa. Biochem J 433: 515–525

Dolinsky TJ, Nielsen JE, McCammon JA, Baker NA (2004) PDB2PQR: an automated pipeline for the setup of Poisson-Boltzmann electrostatics calculations. Nucleic Acids Res 32: W665–667

Forés M, Simon-Carrasco L, Ajuria L, Samper N, Gonzalez-Crespo S, Drosten M, Barbacid M, Jimenez G (2017) A new mode of DNA binding distinguishes Capicua from other HMG-box factors and explains its mutation patterns in cancer. PLoS Genet 13: e1006622

Fryer JD, Yu P, Kang H, Mandel-Brehm C, Carter AN, Crespo-Barreto J, Gao Y, Flora A, Shaw C, Orr HT et al (2011) Exercise and genetic rescue of SCA1 via the transcriptional repressor Capicua. Science 334: 690–693

Futran AS, Kyin S, Shvartsman SY, Link AJ (2015) Mapping the binding interface of ERK and transcriptional repressor Capicua using photocrosslinking. Proc Natl Acad Sci U S A 112: 8590–8595

Gleize V, Alentorn A, Connen de Kerillis L, Labussiere M, Nadaradjane AA, Mundwiller E, Ottolenghi C, Mangesius S, Rahimian A, Ducray F et al (2015) CIC inactivating mutations identify aggressive subset of 1p19q codeleted gliomas. Ann Neurol 78: 355–374

Graham C, Chilton-MacNeill S, Zielenska M, Somers GR (2012) The CIC-DUX4 fusion transcript is present in a subgroup of pediatric primitive round cell sarcomas. Hum Pathol 43: 180–189

Holm L (2020) DALI and the persistence of protein shape. Protein Sci 29: 128–140

Icgc Tcga Pan-Cancer Analysis of Whole Genomes Consortium (2020) Pan-cancer analysis of whole genomes. Nature 578: 82–93

Jimenez G, Guichet A, Ephrussi A, Casanova J (2000) Relief of gene repression by Torso RTK signaling: role of capicua in Drosophila terminal and dorsoventral patterning. Genes & Development 14: 224–231

Jimenez G, Shvartsman SY, Paroush Z (2012) The Capicua repressor--a general sensor of RTK signaling in development and disease. J Cell Sci 125: 1383–1391

Kabsch W (2010) XDS. Acta Crystallographica Section D: Biological Crystallography 66: 125–132

Kamachi Y, Kondoh H (2013) Sox proteins: regulators of cell fate specification and differentiation. Development 140: 4129–4144

Kawamura-Saito M, Yamazaki Y, Kaneko K, Kawaguchi N, Kanda H, Mukai H, Gotoh T, Motoi T, Fukayama M, Aburatani H et al (2006) Fusion between CIC and DUX4 up-regulates PEA3 family genes in Ewing-like sarcomas with t(4;19)(q35;q13) translocation. Hum Mol Genet 15: 2125–2137

Keenan SE, Blythe SA, Marmion RA, Djabrayan NJ, Wieschaus EF, Shvartsman SY (2020) Rapid dynamics of signal-dependent transcriptional repression by capicua. Dev Cell 52: 794–801 e794

Kim JW, Ponce RK, Okimoto RA (2021) Capicua in human cancer. Trends Cancer 7: 77–86

Klaus M, Prokoph N, Girbig M, Wang X, Huang Y-H, Srivastava Y, Hou L, Narasimhan K, Kolatkar PR, Francois M et al (2016) Structure and decoy-mediated inhibition of the SOX18/Prox1-DNA interaction. Nucleic Acids Research 44: 3922–3935

Lam YC, Bowman AB, Jafar-Nejad P, Lim J, Richman R, Fryer JD, Hyun ED, Duvick LA, Orr HT, Botas J et al (2006) ATAXIN-1 interacts with the repressor Capicua in its native complex to cause SCA1 neuropathology. Cell 127: 1335–1347

Lee H, Song JJ (2019) The crystal structure of Capicua HMG -box domain complexed with the ETV 5-DNA and its implications for Capicua-mediated cancers. The FEBS Journal

Lee Y (2020) Regulation and function of capicua in mammals. Exp Mol Med 52: 531–537

Lee Y, Fryer JD, Kang H, Crespo-Barreto J, Bowman AB, Gao Y, Kahle JJ, Hong JS, Kheradmand F, Orr HT et al (2011) ATXN1 protein family and CIC regulate extracellular matrix remodeling and lung alveolarization. Dev Cell 21: 746–757

Liao S, Davoli T, Leng Y, Li MZ, Xu Q, Elledge SJ (2017) A genetic interaction analysis identifies cancer drivers that modify EGFR dependency. Genes Dev 31: 184–196

Liebschner D, Afonine PV, Baker ML, Bunkoczi G, Chen VB, Croll TI, Hintze B, Hung LW, Jain S, McCoy AJ et al (2019) Macromolecular structure determination using X-rays, neutrons and electrons: recent developments in Phenix. Acta Crystallogr D 75: 861–877

Lu C, Wu C, Ghoreishi D, Chen W, Wang L, Damm W, Ross GA, Dahlgren MK, Russell E, Von Bargen CD et al (2021) OPLS4: Improving force field accuracy on challenging regimes of chemical space. J Chem Theory Comput 17: 4291–4300

Lu HC, Tan Q, Rousseaux MW, Wang W, Kim JY, Richman R, Wan YW, Yeh SY, Patel JM, Liu X et al (2017) Disruption of the ATXN1-CIC complex causes a spectrum of neurobehavioral phenotypes in mice and humans. Nat Genet 49: 527–536

Lu M, Yang J, Ren Z, Sabui S, Espejo A, Bedford MT, Jacobson RH, Jeruzalmi D, McMurray JS, Chen X (2009) Crystal Structure of the Three Tandem FF Domains of the Transcription Elongation Regulator CA150. Journal of Molecular Biology 393: 397–408

McCoy AJ, Grosse-Kunstleve RW, Adams PD, Winn MD, Storoni LC, Read RJ (2007) Phaser crystallographic software. J Appl Crystallogr 40: 658–674

Okimoto RA, Breitenbuecher F, Olivas VR, Wu W, Gini B, Hofree M, Asthana S, Hrustanovic G, Flanagan J, Tulpule A et al (2017) Inactivation of Capicua drives cancer metastasis. Nat Genet 49: 87–96

Papagianni A, Fores M, Shao W, He S, Koenecke N, Andreu MJ, Samper N, Paroush Z, Gonzalez-Crespo S, Zeitlinger J et al (2018) Capicua controls Toll/IL-1 signaling targets independently of RTK regulation. Proc Natl Acad Sci U S A 115: 1807–1812

Park S, Lee S, Lee CG, Park GY, Hong H, Lee JS, Kim YM, Lee SB, Hwang D, Choi YS et al (2017) Capicua deficiency induces autoimmunity and promotes follicular helper T cell differentiation via derepression of ETV5. Nat Commun 8: 16037

Paul S, Yang L, Mattingly H, Goyal Y, Shvartsman SY, Veraksa A (2020) Activation-induced substrate engagement in ERK signaling. Mol Biol Cell 31: 235–243

Salam NK, Adzhigirey M, Sherman W, Pearlman DA (2014) Structure-based approach to the prediction of disulfide bonds in proteins. Protein Eng Des Sel 27: 365–374

Sastry GM, Adzhigirey M, Day T, Annabhimoju R, Sherman W (2013) Protein and ligand preparation: parameters, protocols, and influence on virtual screening enrichments. J Comput Aided Mol Des 27: 221–234

Schrodinger L, 2015. The PyMOL Molecular Graphics System, Version 1.8.

Simon-Carrasco L, Grana O, Salmon M, Jacob HKC, Gutierrez A, Jimenez G, Drosten M, Barbacid M (2017) Inactivation of Capicua in adult mice causes T-cell lymphoblastic lymphoma. Genes Dev 31: 1456–1468

Simon-Carrasco L, Jimenez G, Barbacid M, Drosten M (2018) The Capicua tumor suppressor: a gatekeeper of Ras signaling in development and cancer. Cell Cycle 17: 702–711

Tan Q, Brunetti L, Rousseaux MWC, Lu HC, Wan YW, Revelli JP, Liu Z, Goodell MA, Zoghbi HY (2018) Loss of Capicua alters early T cell development and predisposes mice to T cell lymphoblastic leukemia/lymphoma. Proc Natl Acad Sci U S A 115: E1511–E1519

Tate JG, Bamford S, Jubb HC, Sondka Z, Beare DM, Bindal N, Boutselakis H, Cole CG, Creatore C, Dawson E et al (2019) COSMIC: the catalogue of somatic mutations in cancer. Nucleic Acids Research 47: D941–D947

Tseng AS, Tapon N, Kanda H, Cigizoglu S, Edelmann L, Pellock B, White K, Hariharan IK (2007) Capicua regulates cell proliferation downstream of the receptor tyrosine kinase/ras signaling pathway. Curr Biol 17: 728–733

Vivekanandan S, Moovarkumudalvan B, Lescar J, Kolatkar PR Crystal structure of HMG domain of the chondrogenesis master regulator, Sox9 in complex with ChIP-Seq identified DNA element (2015) doi: 10.2210/pdb4S2Q/pdb

Wang B, Krall EB, Aguirre AJ, Kim M, Widlund HR, Doshi MB, Sicinska E, Sulahian R, Goodale A, Cowley GS et al (2017) ATXN1L, CIC, and ETS transcription factors modulate sensitivity to MAPK pathway inhibition. Cell Rep 18: 1543–1557

Williams CJ, Headd JJ, Moriarty NW, Prisant MG, Videau LL, Deis LN, Verma V, Keedy DA, Hintze BJ, Chen VB et al (2018) MolProbity: More and better reference data for improved all-atom structure validation. Protein Science 27: 293–315

Winther JR, Thorpe C (2014) Quantification of thiols and disulfides. Biochim Biophys Acta 1840: 838–846

Zhu K, Day T, Warshaviak D, Murrett C, Friesner R, Pearlman D (2014) Antibody structure determination using a combination of homology modeling, energy-based refinement, and loop prediction. Proteins 82: 1646–1655

