## Supplementary material for "Molecular basis of DNA recognition by the HMG-box-C1 module of Capicua": Electronic Supporting Information Figures and Tables

Supporting Information Table of Contents

1. **Supporting Figures:**

**Figure S1. Related to Figure 1.** The expression construct for $CIC^{min}$

**Figure S2. Related to Figure 1.** $CIC^{min}$ is able to bind the TGAATGAA consensus sequence

**Figure S3. Related to Figure 2.** Representative electron density

**Figure S4. Related to Figure 2.** Sequence numbering alignments between capicua and $CIC^{min}$ (PDB ID 7M5W)

**Figure S5. Related to Figure 2.** Sequence alignments of capicua with representative FF domains

**Figure S6. Related to Figure 3.** The r.m.s.f. values of three MD trajectories of $CIC^{min}$

**Figure S7. Related to Figure 3.** The r.m.s.d. values of $CIC^{min}$ helices from three MD trajectories

**Figure S8. Related to Figures 2 and 3.** Overlayed frames from an MD simulation and overlays of three different MD simulations

**Figure S9. Related to Figure 3.** Building of the $CIC^{min}$ linker at extended C1 domain positions

**Figure S10. Related to Figure 3.** Structures and difference maps of $CIC^{min}$ with and without flipped DNA

**Figure S11. Related to Figure 4.** Overlay of $CIC^{min}$ and the 6JRP structure

**Figure S12. Related to Figures 3 and 4.** Confirmation of the DNA placement in $CIC^{min}$

**Figure S13. Related to Figure 1.** Cosmic database mutations of the capicua C1 domain

2. **Supporting Tables:**

**Table S1. Related to Figure 2.** Nearest structural homologs to the CIC HMG box domain identified from the DALI server.

**Table S2 Related to Figure 2.** Nearest structural homologs to the CIC C1 domain identified from the DALI server.

**Table S3. Related to Figure 2.** Crystallography data and statistics table

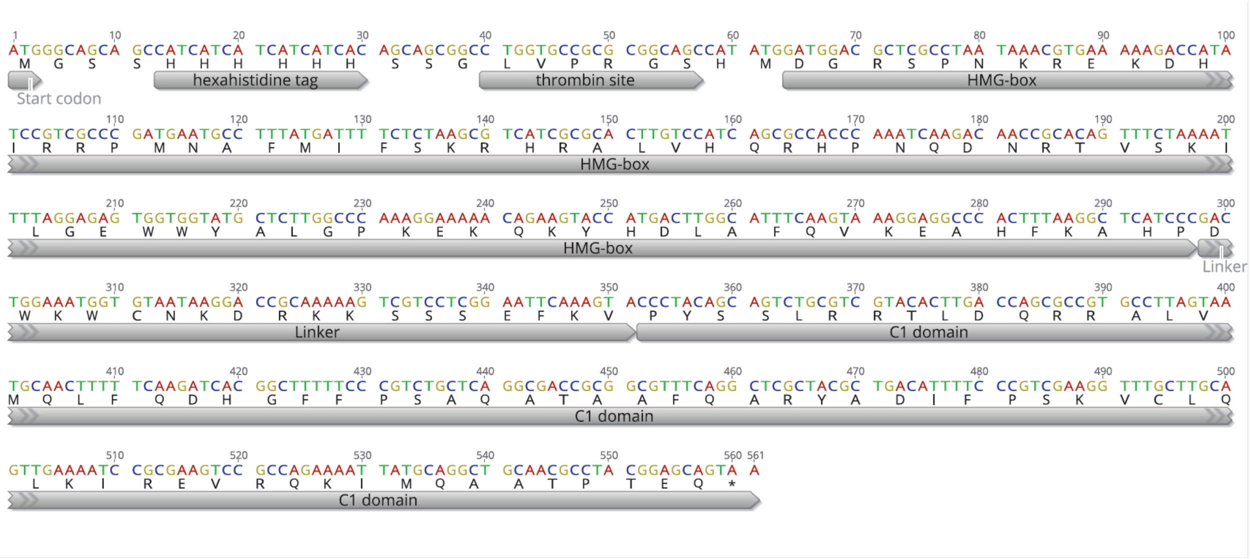


**Figure S1.** Codon optimized sequence, for *E. coli*, encoding the HMG-box and C1 domain of CIC$^{min}$ with human capicua (AF363689_1). The following components of the expression construct are labeled: hexahistidine affinity tag, thrombin cleavage site, HMG-box domain, protein linker region, and C1 domain.

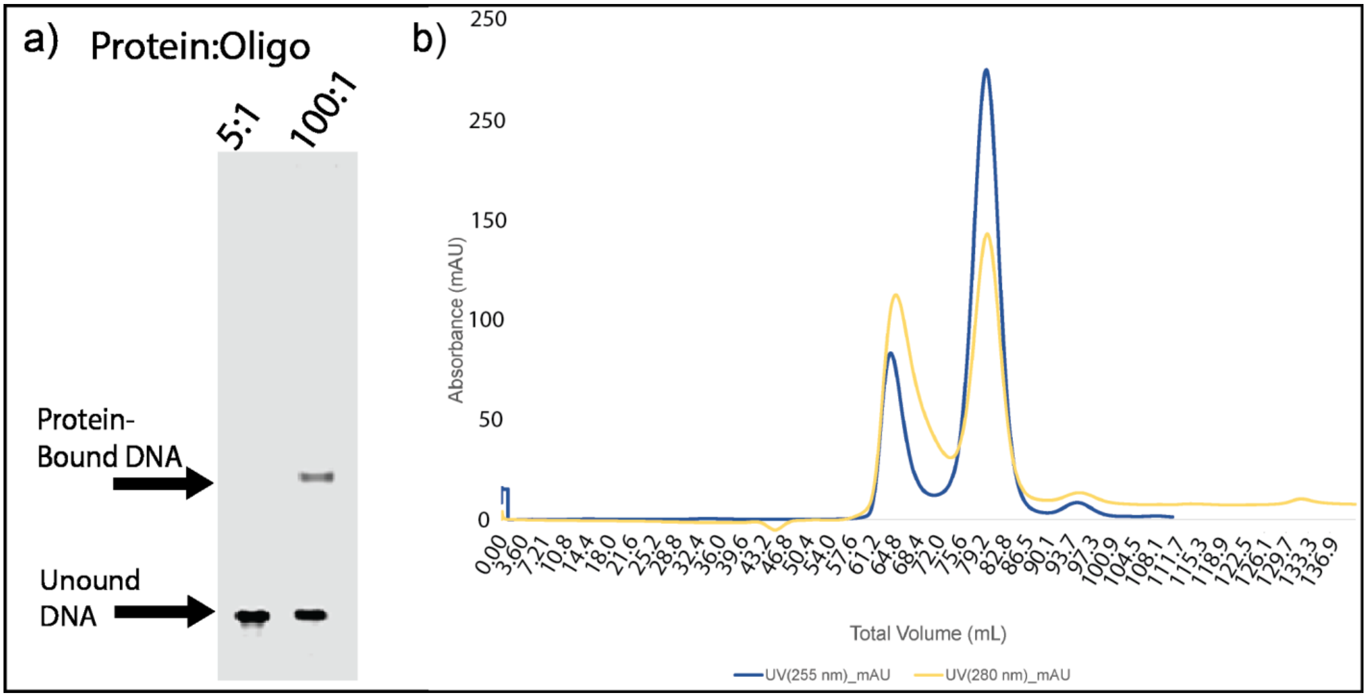


**Figure S2.** EMSA with varying ratios of CIC$^{min}$ to DNA oligomer and size exclusion chromatography (SEC) of CIC$^{min}$ with oligomer shows binding of CIC$^{min}$ with oligomer. **a)** The observed mobility shift for labeled oligo when incubated with CIC$^{min}$ at high protein:oligo ratio supports binding of the oligomer by the CIC$^{min}$ construct. Experiments were performed with 100 pmoles of oligonucleotide. **b)** SEC shows an increased 280 nm absorption over 260 nm for the CIC$^{min}$ DNA complex at approximately 65 mL. This fraction contains the protein and oligomer DNA, and the shoulder at 280 nm is presumed to be unbound protein. The peak at approximately 80 mL is representative of excess, unbound oligo DNA.

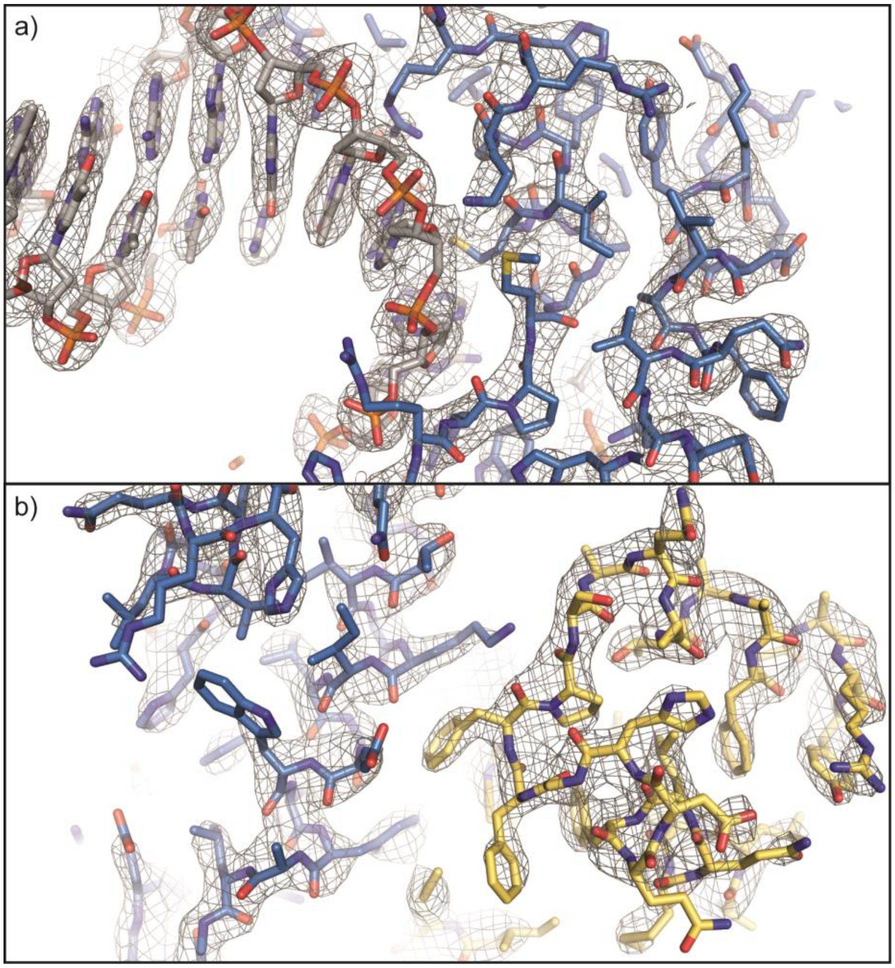


**Figure S3.** Representative electron density figures for the CIC$^{min}$ structure. A composite 2Fo-Fc composite omit electron density map (grey mesh) is displayed for the entire structure at a contour level of 1$\sigma$. Regions of the protein showing electron density for **a)** the DNA and the HMG-box domain and **b)** the HMG domain and C1 domain are displayed. Carbons for the HMG box and C1 domains are colored blue and yellow, respectively, and carbons for DNA are colored grey.

**A**

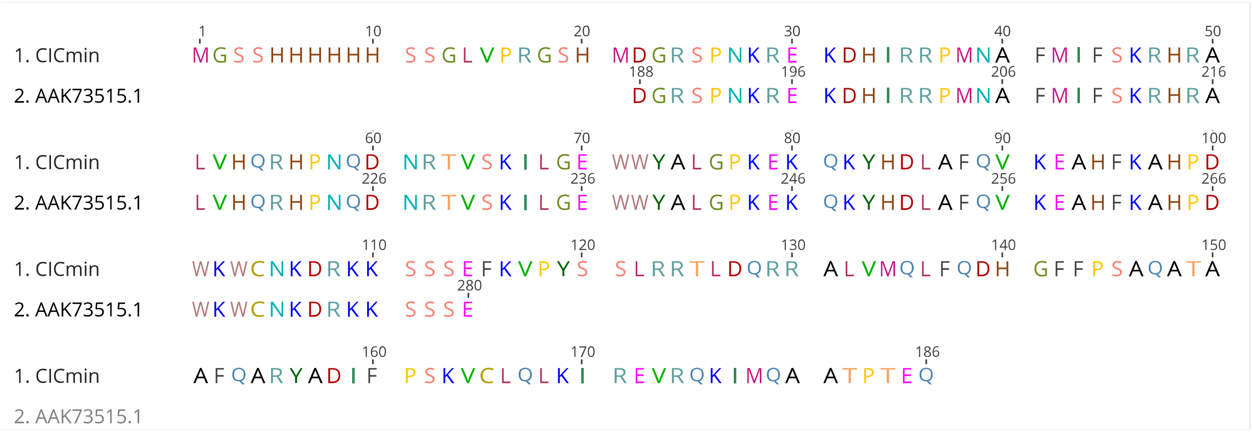


**B**

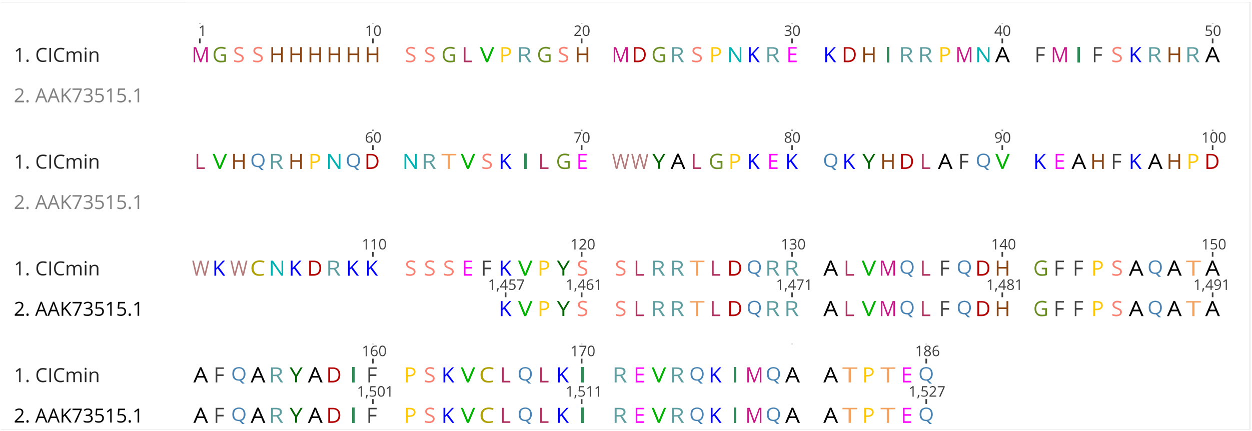


**Figure S4. a)** Sequence alignment of the HMG box domain of $CIC^{min}$ with human capicua (GenBank ID AAK73515.1). **b)** Sequence alignment of the C1 domain of $CIC^{min}$ with human capicua (GenBank ID AAK73515.1).

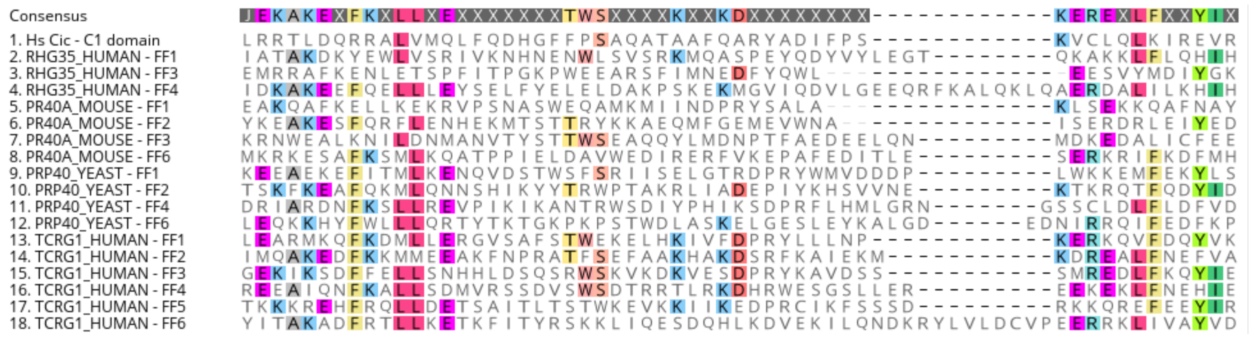

**Figure S5.** Sequence comparison of the capicua C1 domain to representative FF domains. The FF domain is named after two conserved phenylalanine residues present in the first and third helices, which are not present in capicua. Sequence alignment was generated within Geneious Prime.

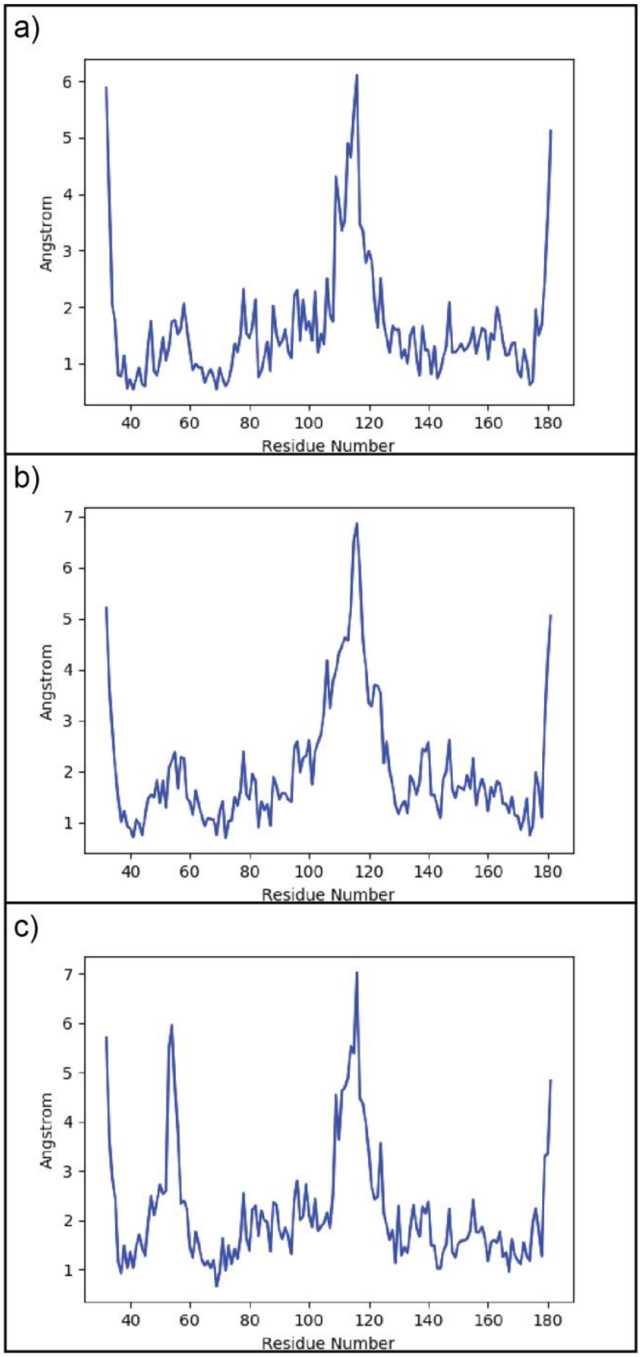


**Figure S6.** The r.m.s.f. values of three trajectories shows the linker and N- and C-terminal regions to consistently be the most disordered of the CIC$^{min}$ structure. **a)** The r.m.s.f. values of a 250 ns trajectory with 100 ps frames shows higher disorder of the loops and linker region of the structure. **b)** The r.m.s.f. values of a 250 ns trajectory with 25 ps frames similarly shows the most disorder in the linker and the N- and C- terminal ends. **c)** The r.m.s.f. values from a 233 ns trajectory with 25 ps frames shows a large increase in the r.m.s.f. values of the C-terminal portion of HMG$^{H1}$, which does not interact with the DNA. This same area of HMG$^{H1}$ shows higher r.m.s.f. values compared to portions of the protein that interact with DNA in the first two trajectories in panels a) and b), but to a lesser extent.

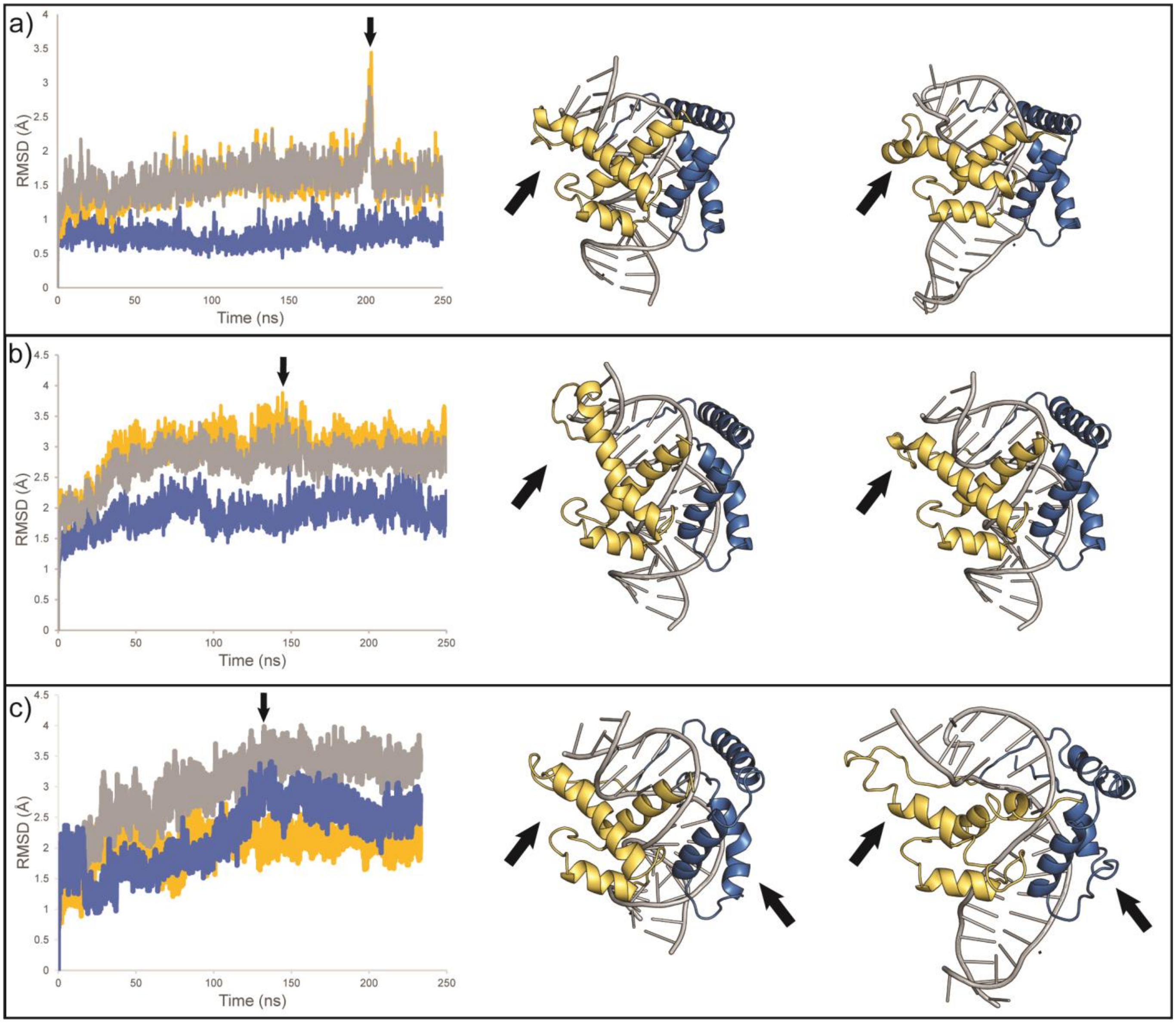


**Figure S7.** The r.m.s.d. values of protein helices from three trajectories indicate the equilibration of the CIC$^{min}$ construct bound to DNA while maintaining the observed binding mode. The HMG-box (blue), C1-domain (yellow), and all protein helices (grey) were each calculated by fitting to the initial frame of the trajectory. Structures shown are the initial state of the trajectory after equilibration (left) and the largest spike in r.m.s.d value (right, with corresponding arrow above the trajectory). Displayed in cartoon form, the HMG-box domain is colored blue, the C1-domain is colored yellow, and the DNA is colored grey. Arrows in each structure indicate regions noted to be more mobile within the HMG-box and the C1-domain during the trajectories. **a)** The r.m.s.d values of the HMG-box and C1-domain of the first trajectory equilibrate around 0.75 Å and 1.5 Å, respectively. **b)** The r.m.s.d. values of the HMG-box and C1-domain of the second trajectory equilibrate around 1.5 Å and 3.0 Å, respectively. **c)** The r.m.s.d. values from a third 233 ns simulation of the HMG-box and C1-domain equilibrate around 2.4 Å and 2.0 Å, respectively. Additionally, the C-terminal end of the $HMG^{H1}$ helix slightly unwinds and thus the r.m.s.d. of the HMG-box is higher than the C1-domain for this simulation. The consensus sequence region of the DNA where protein is bound maintains its shape, whereas the DNA termini are more flexible.

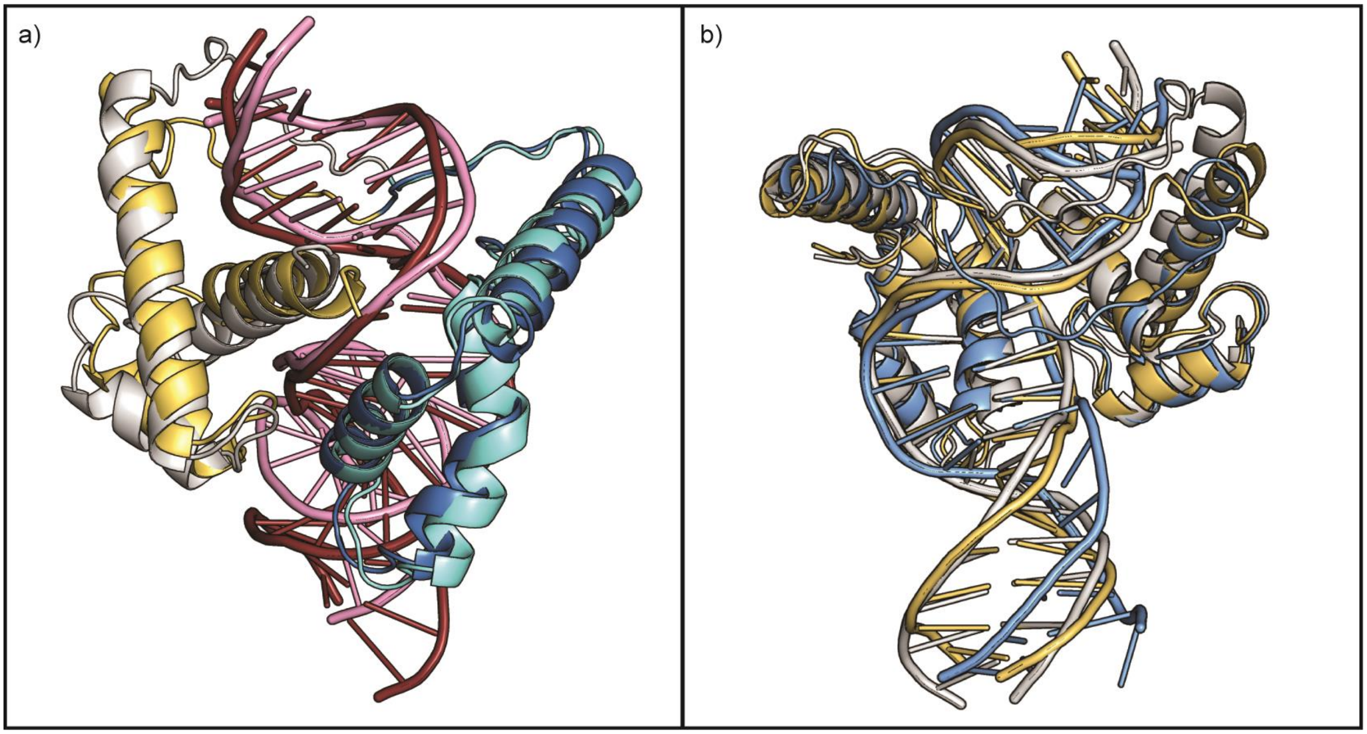


**Figure S8.** The observed DNA binding mode is conserved through three MD simulations, but with variable linker behavior. **a)** An overlay of frames from time 0 and 223.1 ns of a 250 ns MD simulation shows very little change in the overall structure of bound $CIC^{min}$. The HMG-box, C1 domain, and DNA are shown in blue, yellow, and pink, respectively, for the initial frame. For the other frame, they are shown in cyan, white, and red, respectively. **b)** An overlay of states from the three different MD simulations demonstrates three different paths for the interdomain linker. These paths range from branching over the end of the DNA (white) to placing lower along the minor groove (blue).

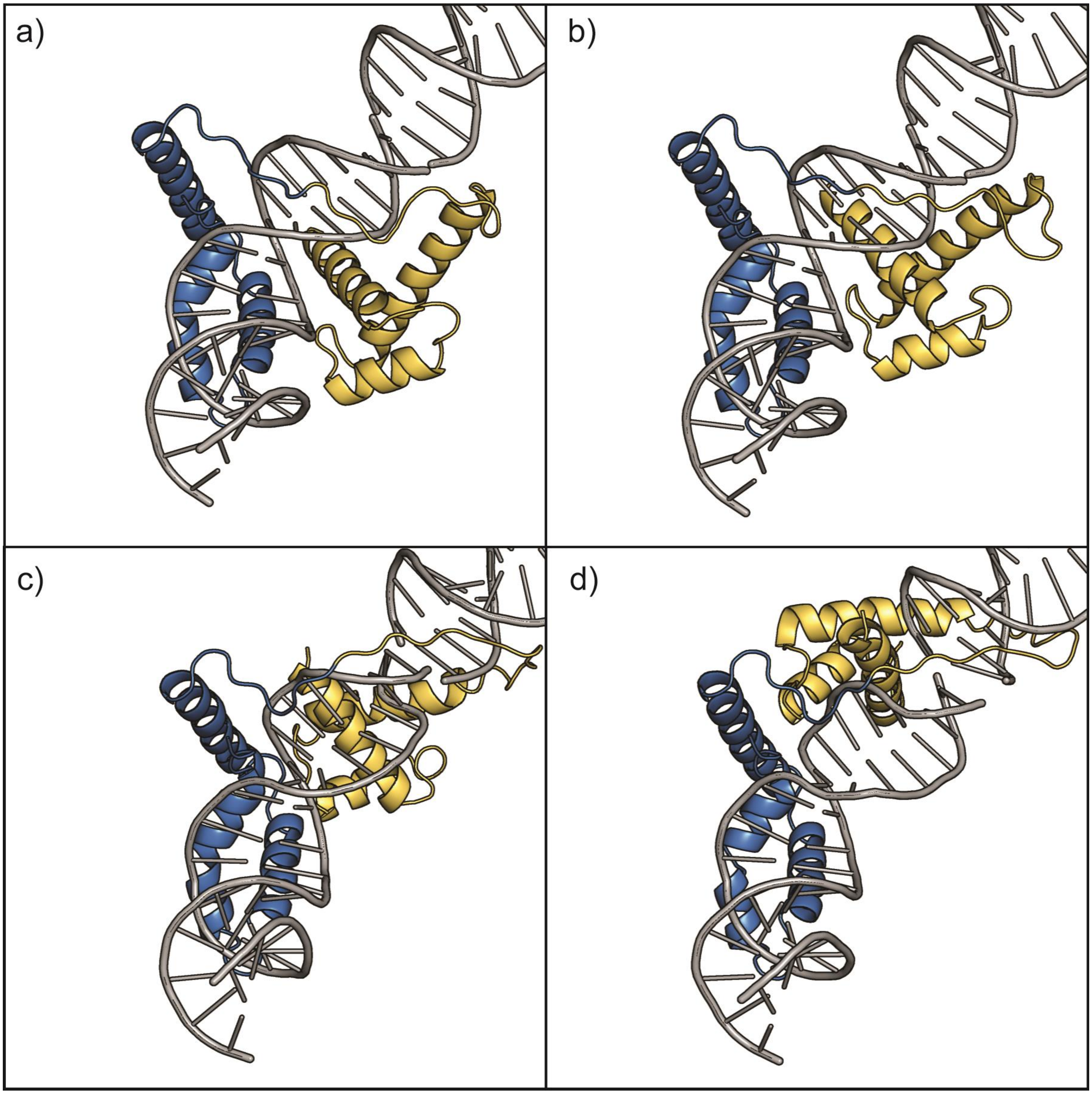


**Figure S9.** The CIC$^{min}$ linker length could accommodate additional movement of the C1 domain. Building of the CIC$^{min}$ linker, using the crosslink proteins function in BioLuminate, was accomplished for structures where the C1 domain was manually moved along the major groove, away from the HMG-box domain. DNA from a symmetry mate was manipulated and moved to artificially extend the DNA. Each successive panel shows the successful building of the linker after further moving away from the HMG-box along the DNA major groove. The automatically built linker wraps around the DNA rather than the C1 domain.

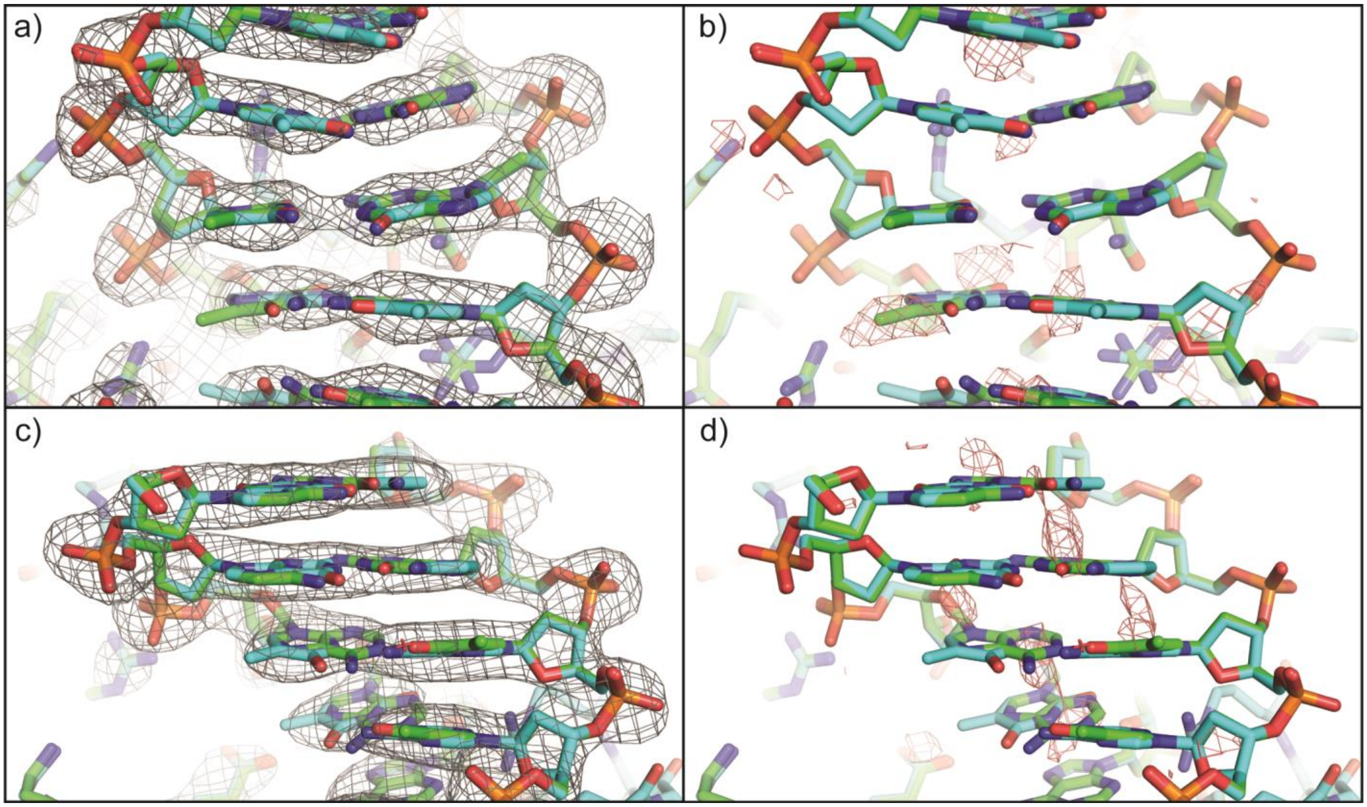


**Figure S10.** The DNA positioning within the CIC$^{min}$-DNA complex was verified through refinement with the DNA flipped. The structure of the proper DNA orientation within CIC$^{min}$ (carbons in blue) is shown aligned with the flipped/incorrectly placed DNA sequence (carbons in green. Panels **a)** and **b)** show the DNA sequence ATGA and panels **c)** and **d)** show the DNA sequence TTTGG (for the properly oriented DNA sequence). Panels **b)** and **d)** show difference Fo-Fc electron density maps resulting from the refinement of DNA in the flipped direction to our model (PDB:7M5W), confirming the flipped DNA orientation is a worse fit to the diffraction data. Positive and negative electron density peaks are shown as green and red mesh, respectively (contoured at a $\pm 3\sigma$ level). The calculated composite omit 2Fo-Fc electron density map is contoured at a $1\sigma$ level and shown in grey mesh.

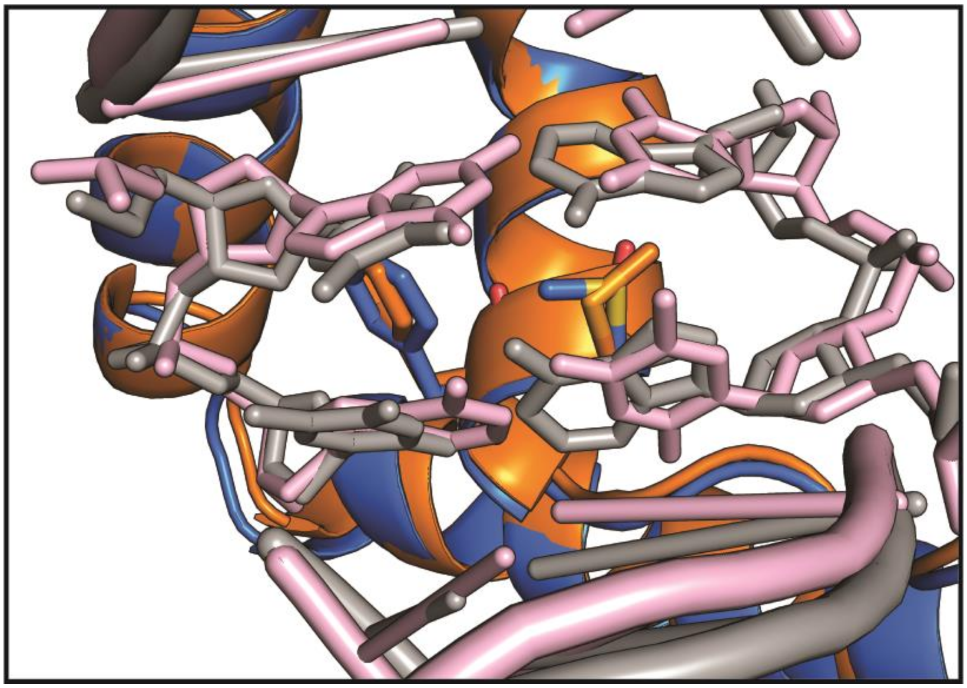

**Figure S11.** The overlay of the CIC$^{min}$-DNA complex and the previously reported CIC HMG-box with DNA (PDB ID 6JRP) shows intercalation at different bases of the consensus sequence. Structures are superimposed based on the HMG-box domain. 6JRP (orange protein and pink DNA) intercalates at the 5´-GA-3´ of the second half of the consensus sequence. CIC$^{min}$ (blue protein and grey DNA) intercalates at the 5´-AA-3´ of the first half of the consensus sequence.

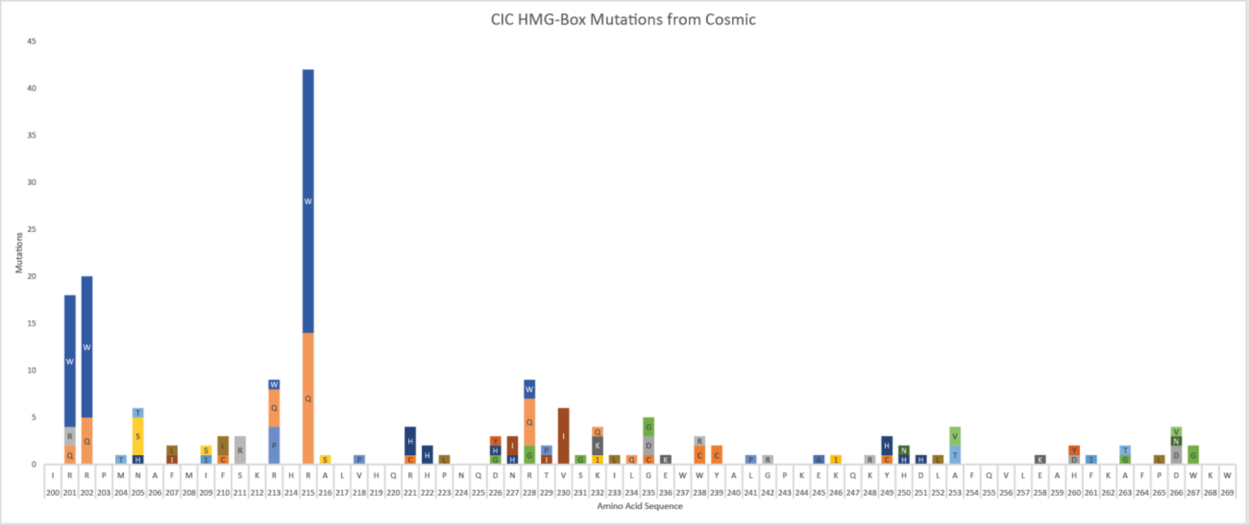


**Figure S12.** Total mutations of residues of the HMG-box reported in the COSMIC database (collected on 05/21/2021).

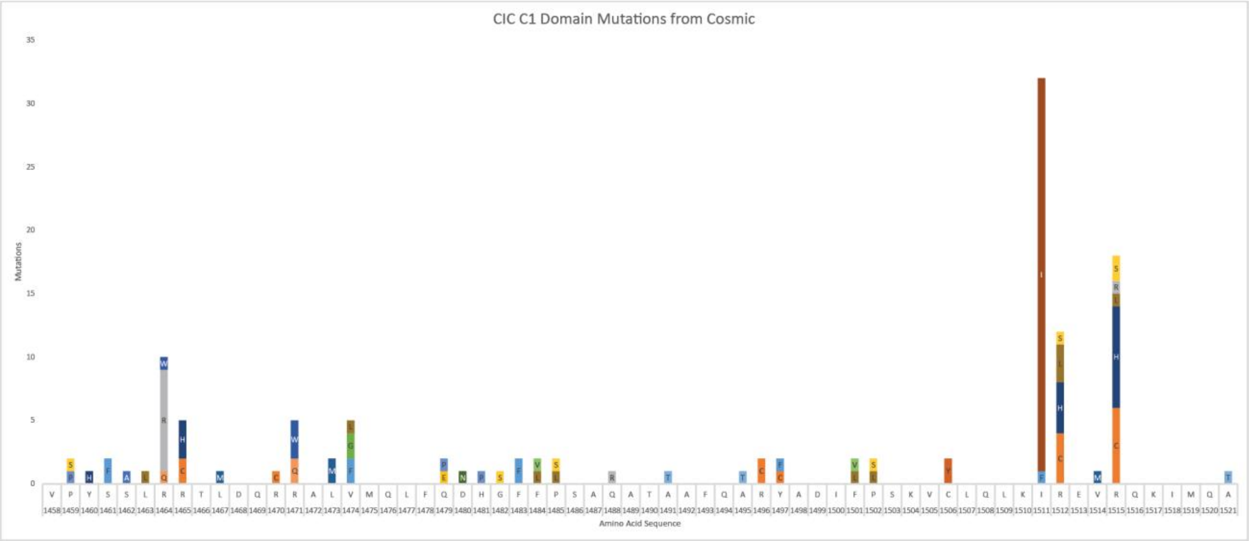


**Figure S13.** Total mutations of residues of the C1-domain reported in the COSMIC database (collected on 05/21/2021).

**Table S1.** Closest structural homologs to the CIC$^{min}$ construct show relatively low sequence identity with other proteins. Nearest structural homologs to the CIC$^{min}$ structure are mainly of the Sox family of transcription factors. Results were collected using the DALI server (Holm, 2020) (12/27/2021).

| Chain | Z | rmsd | lali | %id | PDB Description |
|---|---|---|---|---|---|
| **6jrp** | 12.5 | 0.6 | 73 | 97 | CIC-HMG-ETV5-DNA complex |
| **4y60** | 12 | 1 | 74 | 36 | Sox18-HMG/PROX1-DNA |
| **4s2q** | 11.5 | 1.1 | 74 | 35 | HMG domain of the chondrogenesis master regulator, Sox9 |
| **6l6y** | 11.4 | 1.3 | 74 | 38 | Pluripotency Reprogramming Factor Sox17 mutant (Sox17EK) HMG Domain |
| **3u2b** | 11 | 1.7 | 74 | 34 | Sox4 HMG domain |
| **1j46** | 10.7 | 3 | 78 | 29 | HMG-BOX Domain of the Human Male Sex Determining Factor Sry |
| **5jh0** | 10.6 | 1 | 68 | 22 | mitochondrial DNA packaging protein Abf2p |
| **2lef** | 9.9 | 5.7 | 81 | 32 | Lef1 HMG Domain (From Mouse), Complexed With DNA |
| **2gzk** | 9.5 | 3.5 | 81 | 27 | complex of tandem HMG boxes from HMGB1 and DNA |
| **7cyu** | 9.2 | 1.1 | 62 | 24 | human BAF57 HMG domain |
| **2e6o** | 9.1 | 1.6 | 69 | 38 | HMG box domain from human HMG-box transcription factor 1 |
| **6hb4** | 9 | 5.2 | 81 | 12 | TFAM in Complex with Site-Y |
| **7jjk** | 8.9 | 2.2 | 63 | 32 | Sox30 HMG-domain |
| **2yqi** | 8.8 | 1.8 | 70 | 31 | second HMG-box domain from high mobility group protein B3 |
| **1wz6** | 8.6 | 2.8 | 71 | 52 | HMG-box Domain of Murine Bobby Sox Homolog; Transcription Factor |
| **6cik** | 8.6 | 1.2 | 60 | 35 | Pre-Reaction Complex, RAG1(E962Q)/2-intact/nicked 12/23RSS complex in Mn2+ |
| **6wx8** | 8.5 | 1.7 | 66 | 41 | Importin Subunit Alpha-3 |
| **6l34** | 8.5 | 2.1 | 64 | 23 | HMG domain of human FACT complex subunit SSRP1 |
| **2yul** | 8.4 | 2.5 | 68 | 40 | HMG box of human Transcription factor Sox-17 |

**Table S2.** The top twenty closest structural homologs to the C1 domain show low sequence identity with the closest being Pou3f1. Nearest structural homologs include members of the helix-turn-helix family including the FF domain and the homeodomain. Results were collected using the DALI server (Holm, 2020) (12/27/2021).

| Chain | Z | rmsd | lali | %id | PDB Description |
|---|---|---|---|---|---|
| **3a03** | 6.3 | 1.9 | 52 | 10 | Hox11L1 homeodomain |
| **2h1k** | 6.1 | 2.2 | 54 | 7 | Pdx1 homeodomain in complex with DNA |
| **3nar** | 6 | 2.3 | 56 | 7 | ZHX1 HD4 (zinc-fingers and homeoboxes protein 1, homeodomain 4) |
| **3a01** | 6 | 3.5 | 57 | 11 | Aristaless and Clawless homeodomains bound to DNA |
| **2k85** | 5.9 | 2.3 | 60 | 12 | p190-A RhoGAP FF1 domain |
| **5w66** | 5.9 | 3.1 | 56 | 4 | RNA polymerase I Initial Transcribing Complex State 3 |
| **5hod** | 5.9 | 1.9 | 52 | 10 | LHX4 transcription factor complexed with DNA |
| **5zjr** | 5.9 | 3.2 | 56 | 9 | AbdB/Exd complex bound to a 'Magenta14' DNA sequence |
| **1ig7** | 5.9 | 2.1 | 52 | 12 | Msx-1 Homeodomain/DNA Complex |
| **1d3y** | 5.8 | 2.4 | 54 | 4 | DNA Topoisomerase VI A Subunit |
| **5h7i** | 5.8 | 2.2 | 58 | 7 | Cdt1-MCM2-7 complex in AMPPNP state |
| **4cyc** | 5.8 | 2.5 | 55 | 5 | UBX-EXD-DNA Complex Including the Hexapeptide And Ubda Motifs |
| **1gt0** | 5.8 | 2.1 | 51 | 14 | POU/HMG/DNA ternary complex |
| **6m3d** | 5.8 | 2.7 | 55 | 7 | tandemly connected engrailed homeodomains (EHD) with R53A mutations and DNA complex |
| **5wc9** | 5.7 | 3.4 | 56 | 14 | Human Pit-1 and 4xCATT DNA complex |
| **2xsd** | 5.7 | 2.5 | 53 | 15 | dimeric Oct-6 (Pou3f1) POU domain bound to palindromic MORE DNA |
| **6es3** | 5.7 | 3.2 | 58 | 12 | CDX2-DNA(TCG) |
| **4uus** | 5.7 | 3.1 | 56 | 5 | UBX-EXD-DNA Complex Including the Ubda Motif |
| **1ahd** | 5.7 | 2.5 | 58 | 7 | Antennapedia Homeodomain-DNA Complex |

**Table S3.** Crystallographic table.

| | *CIC^min^ complexed with DNA* |
|---|---|
| Resolution range | 23.45 - 2.95 (3.13 - 2.95) * |
| Space group | $P\ 2_1\ 2_1\ 2_1$ |
| Unit cell | 46.892 79.915 106.963 90 90 90 |
| Total reflections | 79037(12044) |
| Unique reflections | 8838 (1349) |
| Multiplicity | 8.9 (6.6) |
| Completeness (%) | 98.8 (97.3) |
| Mean I/sigma(I) | 22.44 (3.66) |
| Wilson B-factor | 49.76 |
| *R*-sym # | 0.083 (0.506) |
| *CC*1/2 & | 0.999 (0.970) |
| **Refinement** | |
| Reflections used in refinement | 8769 (2705) |
| Resolution | 23.84 – 2.95 (3.38 – 2.95) |
| R-work $ | 0.2093 (0.2471) |
| R-free $ | 0.2434 (0.3135) |
| Number of non-hydrogen atoms | 1912 |
| macromolecules | 1896 |
| metal ion | 1 |
| solvent | 15 |
| RMS(bonds) | 0.003 |
| RMS(angles) | 0.46 |
| Ramachandran favored (%) † | 98.51 |
| Ramachandran allowed (%) | 1.49 |
| Ramachandran outliers (%) | 0 |
| Rotamer outliers (%) | 0.83 |
| Clashscore | 1.15 |
| Average B-factor (Â$^2$) | 67.44 |
| macromolecules | 67.42 |
| ligands | 92.74 |
| solvent | 49.27 |
| *Number of TLS groups* | *4* |

* Highest resolution shell is shown in parentheses

\# $R_{sym} = \Sigma_{hkl}\Sigma_i|I_i(hkl) - \langle I(hkl)\rangle|/\Sigma_{hkl}\Sigma_i I_i(hkl)$.

& $CC^* = [2CC_{1/2}/(1+CC_{1/2})]^{1/2}$ where $CC_{1/2}$ is the correlation between two random half datasets containing half of the measured intensities for each unique reflection and CC* approximates the correlation coefficient for a noise-free dataset.

$ $R_{work} = \Sigma|F_{obs}(hkl) - F_{calc}(hkl)|/\Sigma|F_{obs}(hkl)|$, where $F_{obs}(hkl)$ and $F_{calc}(hkl)$ are the observed and calculated structure factor amplitudes of ~95% of the reflections used for refinement. $R_{free}$ was calculated from the ~5% of total reflections that were omitted from the refinement.

† Ramachandran percentages generated using MolProbity (Williams *et al*, 2018).

**References**

Holm L (2020) DALI and the persistence of protein shape. *Protein Sci* 29: 128-140

Williams CJ, Headd JJ, Moriarty NW, Prisant MG, Videau LL, Deis LN, Verma V, Keedy DA, Hintze BJ, Chen VB *et al* (2018) MolProbity: More and better reference data for improved all-atom structure validation. *Protein Science* 27: 293-315
